# MTCH2 is a mitochondrial outer membrane protein insertase

**DOI:** 10.1101/2022.09.15.508165

**Authors:** Alina Guna, Taylor A. Stevens, Alison J. Inglis, Joseph M. Replogle, Theodore K. Esantsi, Gayathri Muthukumar, Kelly C.L. Shaffer, Maxine L. Wang, Angela N. Pogson, Jeff J. Jones, Brett Lomenick, Tsui-Fen Chou, Jonathan S. Weissman, Rebecca M. Voorhees

**Affiliations:** Whitehead Institute for Biomedical Research, Massachusetts Institute of Technology, Cambridge, MA 02142, USA; Division of Biology and Biological Engineering, California Institute of Technology, 1200 E. California Ave., Pasadena, CA 91125, USA; Medical Scientist Training Program, University of California, San Francisco, San Francisco, CA 4158, USA; Tetrad Graduate Program, University of California, San Francisco, San Francisco, CA 94158, USA; Howard Hughes Medical Institute, Massachusetts Institute of Technology, Cambridge, MA 02142, USA; Department of Biology, Massachusetts Institute of Technology, Cambridge, MA 02142, USA; David H. Koch Institute for Integrative Cancer Research, Massachusetts Institute of Technology, Cambridge, MA,02142, USA

## Abstract

In the mitochondrial outer membrane, tail-anchored (TA) proteins play critical roles in cytoplasmic-mitochondrial communication. Using genome-wide CRISPRi screens, we identify factors involved in mitochondrial TA biogenesis in human cells. We show that MTCH2, and its paralog MTCH1, are required for insertion of biophysically diverse mitochondrial TAs, but not outer membrane β-barrel proteins. In a reconstituted system, purified MTCH2 is sufficient to mediate insertion into proteoliposomes. Functional and mutational studies reveal that MTCH2 uses membrane-embedded hydrophilic residues to function as a gatekeeper for outer membrane protein biogenesis, controlling mislocalization of TAs into the endoplasmic reticulum and the sensitivity of leukemia cells to apoptosis. Our identification of MTCH2 as an insertase provides a mechanistic explanation for the diverse phenotypes and disease states associated with MTCH2 dysfunction.

**One-Sentence Summary:** MTCH2 is both necessary and sufficient for insertion of diverse α-helical proteins into the mitochondrial outer membrane, and is the defining member of a family of insertases that have co-opted the SLC25 transporter fold.

## Main text

Mitochondria are essential organelles of endosymbiotic origin that have evolved to play a central role in eukaryotic cell metabolism and signaling (1, 2). While the regulated fission, fusion, and turnover of mitochondria are critical to cellular viability, they also provide a platform for mediating cellular processes such as apoptosis and the innate immune response. Both mitochondrial function and their ability to communicate with the cytosol depend on proteins embedded in the outer mitochondrial membrane. As a result, dysregulation of outer membrane protein function is associated with ageing and has been implicated in the pathogenesis of a variety of human diseases including Alzheimer’s, Parkinson’s, and many cancers (3–5).

In mammals, the mitochondrial outer membrane contains ∼150 different proteins, all of which are encoded in the nuclear genome and must be targeted and inserted into the membrane. For outer membrane β-barrel proteins, which would have been present in the bacterial predecessor of the mitochondria, this is a two-step process requiring translocation into the intermembrane space (IMS) via the translocase of the outer membrane (TOM) and subsequent insertion by the sorting and assembly machinery (SAM) complex. The SAM complex is evolutionarily derived from the bacterial β-barrel assembly machinery (BAM), which plays an analogous role in prokaryotes (6).

By contrast, how α-helical proteins are inserted into the mitochondrial outer membrane, a function that would not have been present in the original endosymbiont, remains poorly understood (7). In yeast the mitochondrial import protein 1 (Mim1) has been implicated in this process (8). However, the size and complexity of the outer membrane proteome has expanded substantially between fungi and mammals, suggesting metazoans rely on distinct and functionally more complex biogenesis machinery.

One important class of outer membrane proteins are tail-anchored proteins (TAs), which are characterized by a single C-terminal α-helical transmembrane domain (TMD). TA proteins’ soluble N-termini are localized to the cytosol and mediate a highly diverse array of functions including regulating apoptosis (BCL-2 family), the innate immunity (MAVS), and several facets of mitochondrial turnover and dynamics (FIS1, RHOT1/2, BNIP3). Mammalian cells contain ∼45 TAs that must be post-translationally targeted and inserted from their site of synthesis in the cytosol into the outer mitochondrial membrane (9). Numerous conflicting biogenesis pathways have been proposed for mitochondrial TAs. Studies have suggested that yeast mitochondrial TAs may be autonomously inserted, based on the observation that trypsin treated cells remain insertion-competent (10–12). However, other work argues that TA insertion relies on a pathway that can be saturated, implying a protein mediated process, but without direct evidence for the identity of the putative insertase machinery (10). We therefore set-out to systematically identify and characterize the factors required for mitochondrial TA biogenesis in human cells.

Because the TOM complex forms the canonical insertase into mitochondria, and had been previously implicated in TA biogenesis, we first sought to test if it was required for TA insertion. Using an in vitro competition assay with mitochondrially enriched membranes, we confirmed that mitochondrial TAs do not rely on the TOM-mediated pathway for insertion into the outer membrane, consistent with earlier studies (Fig. 1A and fig. S1) (13). To systematically identify factors required for biogenesis of mitochondrial TAs, we adapted a split-GFP reporter system to enable CRISPR-based screens. We constitutively expressed the first 10 β-strands of GFP (GFP1-10) in the inner membrane space and appended the 11th β-strand to the C-terminus of a mitochondrial TA. Successful integration of the TA into the outer membrane results in complementation of the complete GFP (GFP1-10+GFP11) and fluorescence (Fig. 1B and fig. S2) (14). Critically, this strategy specifically monitors substrate integration into the outer mitochondrial membrane, rather than mislocalization to other organelles or the cytosol (e.g., no complementation is observed between a GFP11-tagged ER TA protein and mitochondrial localized GFP1-10; fig. S2B and fig. S2C). We chose the mitochondrial TA OMP25 as a screening substrate because, although OMP25 can be mislocalized to the ER, it is not essential; is monomeric, and thus does not depend on successful assembly for stability; is not cytotoxic when overexpressed; and tolerates a GFP11 tag on its C-terminus that can robustly complement an IMS localized GFP1-10.

**Fig. 1.**
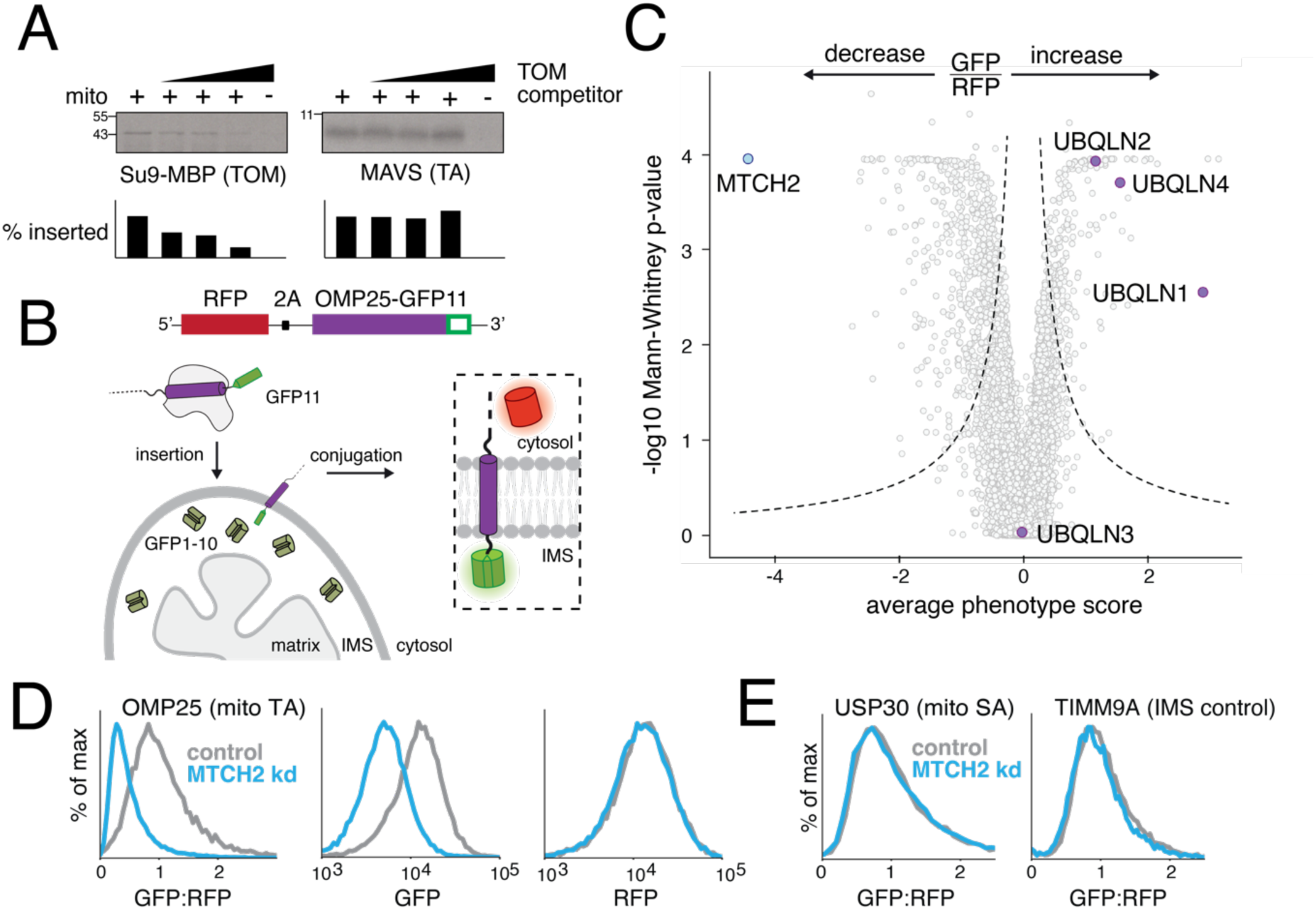
Systematic characterization of human mitochondrial TA biogenesis. **(A)** An ^35^S-methionine labelled TOM substrate (made from a fusion of the canonical TOM targeting sequence Su9 and the globular protein MBP) or MAVS (a mitochondrial TA protein) were translated in rabbit reticulocyte lysate and released from the ribosome using puromycin. Competition assays were performed by incubation with mitochondrially enriched membranes (see fig. S1) in the presence of increasing concentrations of a recombinant TOM competitor (Su9-DHFR). Mitochondrial insertion was assessed by protease protection and analyzed by SDS-PAGE and autoradiography. **(B)** Schematic of the split GFP reporter system used to specifically query integration of substrates into the outer mitochondrial membrane. A mitochondrial membrane protein fused to GFP11 is expressed in a cell constitutively expressing GFP1-10 in the inner membrane space (IMS) along with a translation normalization marker (mCherry, though referred to as RFP throughout for simplicity). Successful integration into the outer membrane results in complementation and GFP fluorescence. **(C)** Volcano plot of GFP:RFP stabilization phenotype [log_2_(High GFP:RFP/low GFP:RFP)] for the three strongest sgRNAs versus Mann-Whitney p values from two independent replicates of the genome-wide CRISPRi screen using OMP25-GFP11. Individual genes are displayed in grey, and specific factors that increase or decrease OMP25 mitochondrial integration are highlighted and labelled. **(D)** Integration into mitochondria of the OMP25-GFP11 reporter described in (B) was assessed in K562 cells expressing a non-targeting (nt) or MTCH2 knock down sgRNA (kd). GFP fluorescence relative to a normalization marker was determined by flow cytometry and displayed as a histogram. **(E)** Biogenesis of USP30-GFP11, an outer membrane resident signal anchored protein, and TIM9A-GFP11, an IMS localized protein, were assessed as in (D).

We engineered a human K562 cell line to stably express an IMS localized GFP1-10 and the CRISPR inhibition (CRISPRi) machinery, which enables programmed repression of genes targeted by a single-guide RNA (sgRNA). We subsequently introduced the full-length endogenous sequence of OMP25 appended to GFP11 (OMP25-GFP11) along with an expression normalization marker (RFP) separated by a viral 2A sequence. By expressing both the TA and RFP from a single open reading frame the resulting GFP:RFP ratio reports on the efficiency of post-translational integration into mitochondria (Fig. 1B). After transduction with a genome-wide CRISPRi sgRNA library, cells with altered TA integration as evidenced by a change in the GFP:RFP fluorescence were isolated using FACS and their associated sgRNAs were identified by deep sequencing (fig. S2D; Table S1) (15). We hypothesized that by using this ratiometric approach, sgRNAs enriched in either the high or low populations of GFP:RFP fluorescence would include factors that affect the post-translational stability or targeting of our reporter, and therefore represent putative quality control or biogenesis factors, respectively.

Indeed, we found that amongst the factors whose depletion led to increased OMP25-GFP fluorescence were the paralogous chaperones ubiquilin-1, -2, and -4 (UBQLNs) (Fig. 1C and fig. S3). UBQLN3 has testis-specific expression, and was therefore not a significant hit under the screening conditions. Increased integration of OMP25 into mitochondria in the absence of the UBQLNs is consistent with their previously reported role in recognizing and degrading mislocalized mitochondrial TAs (16). Similarly, knockdown of the ER membrane protein complex (EMC), which we found was the major pathway for insertion of mislocalized mitochondrial TAs into the ER, also increases insertion into the mitochondria in agreement with recent studies (fig. S4) (17). Presumably, depletion of either the UBQLNs or the EMC increases the cytosolic pool of OMP25, allowing additional opportunities for successful insertion into the outer membrane prior to degradation by the ubiquitin proteasome pathway.

Conversely, depletion of the outer membrane resident mitochondrial carrier homologue 2 (MTCH2) resulted in by far the most pronounced loss of OMP25 integration. Validation with a single guide knockdown confirmed that loss of MTCH2 specifically decreases GFP fluorescence without affecting the levels of our soluble expression control (Fig. 1D), consistent with a potential role in targeting or insertion of OMP25-GFP11 into mitochondria. These effects were independent of the IMS-targeting sequence used for the GFP1-10, suggesting that the observed MTCH2 phenotype is not mediated by changes to GFP1-10 levels (fig. S5A). MTCH2 is a member of the solute carrier 25 (SLC25) family, integral membrane proteins best known for their role in transporting metabolites into the mitochondrial matrix, but its sequence suggests its function has potentially diverged, and it has no known substrates or transporter activity (18). Further, loss of MTCH2 is associated with a variety of pleotropic phenotypes including defects in mitochondrial fusion, lipid homeostasis, and apoptosis (19–21). However, the underlying biochemical activity of MTCH2 is not known.

Because of the diverse phenotypes attributed to MTCH2, we sought to exclude the possibility that general dysregulation of mitochondria, the outer membrane, or lipogenesis, could explain the observed biogenesis defect on OMP25. (i) We found that factors involved in mitochondrial lipid biosynthesis, some of which are required for MTCH2-dependent mitochondrial fusion, were not significant hits in our reporter screen (fig. S5B). (ii) Additionally, treatment of cells with the pan GPAT inhibitor FSG67, which phenocopies the effect of MTCH2 depletion on mitochondrial fission by downregulating the synthesis of lysophosphatidic acid, did not affect our OMP25 reporter (fig. S5C) (19). This result suggests that the effect of MTCH2 on TA biogenesis cannot be solely explained by changes to the lipid composition of the outer membrane. (iii) Further, a fluorescent reporter for the outer membrane protein USP30 was unaffected by MTCH2 depletion, excluding general dysregulation of the outer membrane upon loss of MTCH2 (Fig. 1E). (iv), Finally, we confirmed that loss of MTCH2 did not affect the biogenesis of an IMS resident protein, TIM9A, discounting general defects on mitochondrial insertion (Fig. 1E).

We next sought to determine if MTCH2 could be playing a more general role in the biogenesis of other mitochondrial outer membrane proteins. Using a quantitative proteomics strategy, we compared the steady-state levels of proteins in mitochondria purified from wildtype or MTCH2 depleted cells (Fig. 2A; Table S2). We identified several endogenous α-helical proteins localized to the outer membrane that were reproducibly depleted upon loss of MTCH2. These included TA (RHOT2, CYB5B), signal anchored (HSDL1, TDRKH), and several multipass proteins (FUNDC1, PLGRKT), a subset of which we confirmed by western blotting (Fig. 2B and fig. S6). By contrast, MTCH2 depletion does not affect the steady state levels of VDAC1, a highly abundant β-barrel protein on the outer membrane, and CYC1, an inner membrane α-helical protein. Because loss of MTCH2 does not appreciably alter mRNA levels across the transcriptome, we concluded that the effects of MTCH2 on the mitochondrial outer membrane proteome must be occurring post-transcriptionally (22).

**Fig. 2.**
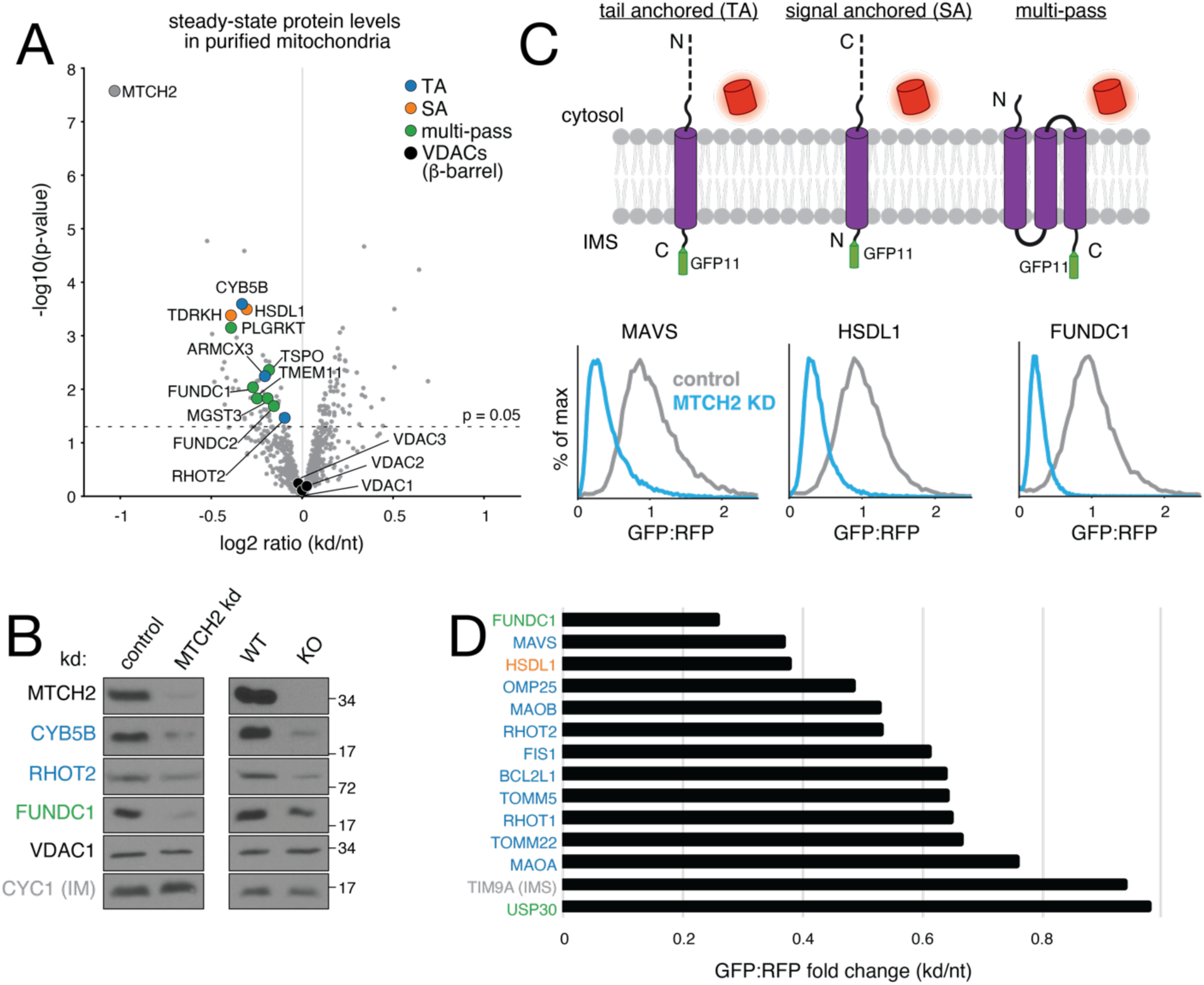
MTCH2 is required for mitochondrial outer membrane protein biogenesis. **(A)** Label-free mass spectrometry analysis of purified mitochondria isolated from K562 cells using a percoll gradient (fig. S1B) expressing a MTCH2 targeting sgRNA (kd) compared to a non-targeting control (nt). Displayed are proteins that across four biological replicates were statistically altered in MTCH2 depleted versus non-targeting guide expressing cells. **(B)** Western blotting of endogenous proteins from MTCH2 depleted (kd) and control cells in (generated as in A; left), and wild type and MTCH2 knock out (KO) cells (right). Substrates are colored by topology based on the key shown in (B). **(C)** Flow cytometry analysis of integration of outer membrane protein reporters using the split GFP system described in Fig. 1B. GFP fluorescence relative to an RFP expression control are displayed as histograms in MTCH2 knockdown versus non-targeting K562 CRISPRi cells. Displayed are representative examples of a TA, SA, and multipass membrane protein that have a MTCH2 dependent biogenesis defect. **(D)** Summary of dependence on MTCH2 for the indicated outer membrane substrates determined using the fluorescent reporter system shown in (C) and colored by topology based on the key in (B).

To test whether MTCH2 is functioning directly in biogenesis, we verified several of these hits, as well as a panel of mitochondrial TA proteins, using our ratiometric fluorescent reporter strategy. For these experiments, we used the full-length endogenous sequences of all substrates tested. We found that both knockdown (Fig. 2C and Fig. 2D) and knockout (fig. S7) of MTCH2 reproducibly affected the biogenesis of a functionally diverse set of TA proteins that contain TMDs with varying hydrophobicity, flanking charge, and helical propensity. We further verified that at least one multipass protein, which was de-enriched in our proteomics (FUNDC1), displays a MTCH2 dependent biogenesis defect. FUNDC1’s third TMD adopts a similar topology to a TA protein, which may partially explain MTCH2’s broader role in outer membrane protein biogenesis beyond TAs (23).

Based on these experiments, its localization, and its evolutionary divergence from the transporter family, we reasoned that MTCH2 may have evolved the ability to insert α-helical proteins into the outer membrane. To test this hypothesis, we focused on TA proteins, because they are the largest class of α-helical outer membrane proteins and adopt a uniform topology, facilitating in vitro study. Therefore, to delineate the role of MTCH2 in TA biogenesis, we employed an in vitro insertion and protease protection assay (fig. S1D). We translated the full-length endogenous OMP25 in the mammalian rabbit reticulocyte lysate (RRL) system. The resulting nascent substrates were incubated with mitochondria isolated from either wild type or MTCH2 depleted cells, and treated with protease to degrade all but those regions protected by insertion into/across the membrane. Because proteasomal degradation is inactive in RRL, loss of a protected fragment necessarily reflects an effect at the level of insertion and not a change to protein quality control, providing additional information beyond our cell-based reporter assay. We observed a protected fragment consistent in size to the TMD and C-terminus of OMP25 that is partially lost in the absence of MTCH2 (fig. S8C; Fig. 3A). This effect is dependent on the TMD, as fusions of the OMP25 TMD with unrelated N-terminal domains recapitulated the requirement for MTCH2 for insertion (fig. S8D).

**Fig. 3.**
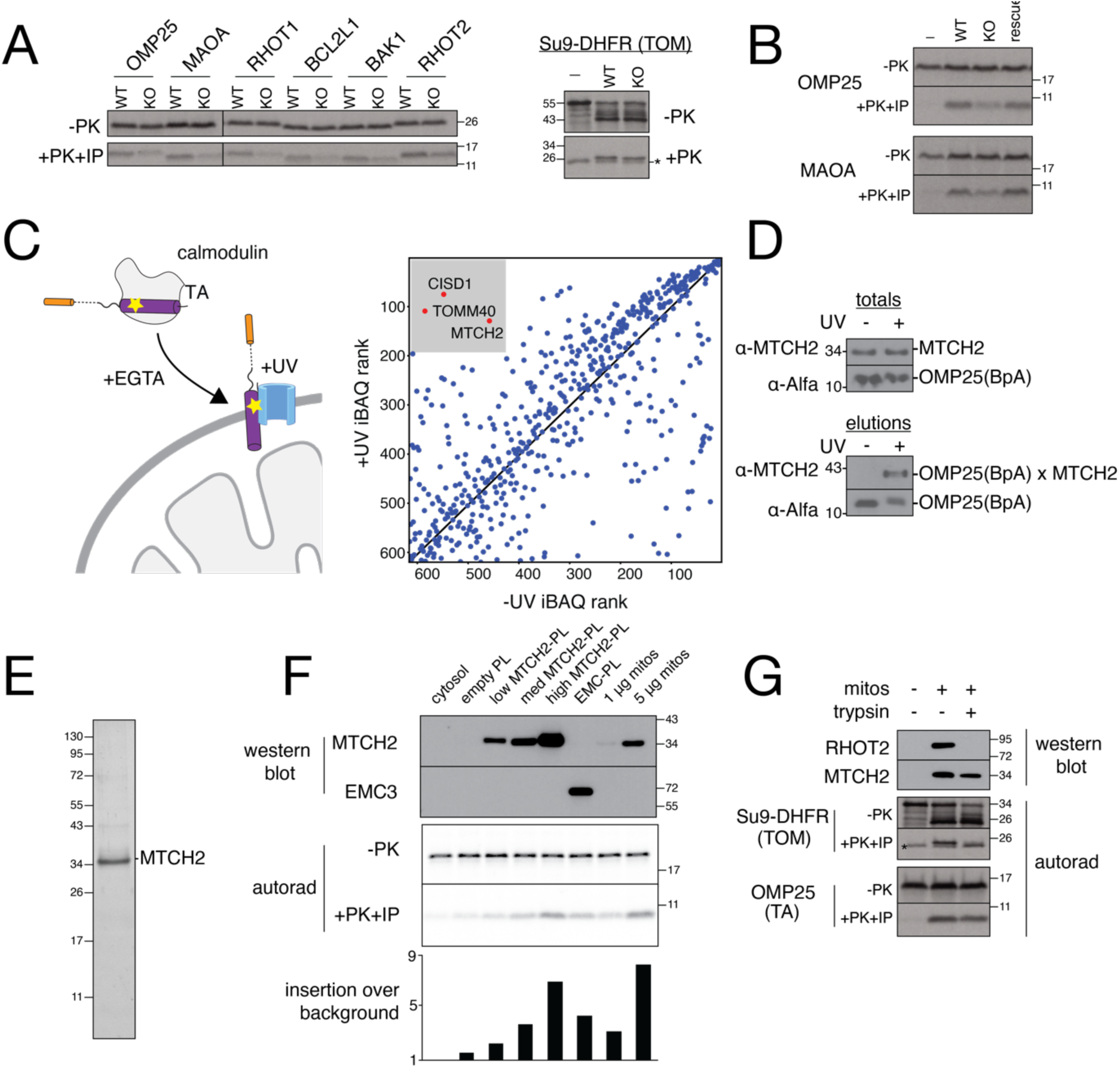
MTCH2 inserts diverse mitochondrial TAs into the outer membrane. **(A)** The indicated ^35^S-methionine labelled TA proteins (fig. S8B) were analyzed for in vitro insertion into mitochondria isolated from wild type (WT) or MTCH2 knockout (KO) K562 cells. Displayed are the samples prior to addition of protease (-PK) and the protease protected fragment that has been affinity purified via a 6xHIS tag on the C-terminus of each substrate (PK+IP), ensuring insertion in the correct topology. Canonical TOM substrate controls that are targeted to either the matrix (Su9-DHFR) or IMS (cyt. b_2_-DHFR) were tested in parallel and are depicted on the right and in fig. S9C. *Denotes the folded, protease resistant, DHFR domain that migrates immediately below the mature, matrix targeted control, and is visible in the absence of mitochondria. **(B)** As in (A) comparing insertion of the indicated TA proteins into wild type, MTCH2 KO, and MTCH2 KO + MTCH2 rescue mitochondria. **(C)** (left) Schematic showing the photocrosslinking strategy. OMP25 containing the photoactivatable amino acid BpA within its TMD was expressed and purified from *E. coli* as a complex with calmodulin. OMP25^BpA^ was released from calmodulin by addition of EGTA in the presence of mitochondria purified from K562 cells using a percoll gradient (fig. S1B). Crosslinking was activated by UV-irradiation, and the resulting crosslinked species were affinity purified via the Alfa-tag on the N-terminus of OMP25^BpA^ for identification by mass spectrometry. (right) All proteins identified by mass spectrometry were ranked by iBAQ abundance, and those specifically enriched in the UV compared to the -UV control are highlighted. **(D)** As in (C) with the resulting elution analyzed by western blotting to assess levels of crosslinked OMP25 ^BpA^-MTCH2. (**E)** MTCH2 was expressed and purified from human cells and analyzed by SDS-PAGE and Sypro-Ruby staining. **(F)** Following reconstitution, the recovered proteoliposomes were analyzed by western blotting for incorporation of the indicated proteins in comparison to purified mitochondria. Using a protease protection assay, OMP25 synthesized in native cytosol was tested for insertion into liposomes reconstituted with increasing amounts of purified MTCH2, the human EMC, or an empty control. In parallel for comparison, we tested insertion into mitochondria isolated from K562s at two concentrations. The resulting protease protected fragments were immunoprecipitated, imaged by autoradiography (autorad), and their intensity quantified on a phosphoimager, and are displayed normalized to the cytosol only control. The presence of a small amount of protected fragment in cytosol alone may be indicative of chaperone binding. **(G)** Mitochondria from WT K562 cells were treated with trypsin and their ability to insert TOM substrates (Su9-DHFR) and TA substrates (OMP25) was assayed by protease protection as in (A). The OM protein RHOT2 was confirmed to be degraded in a trypsin-dependent manner by western blot, while MTCH2 remained largely intact.

To test whether TMDs and their flanking regions confer differential dependence on MTCH2, we generated a suite of fusion proteins containing the C-terminal TMDs of several mitochondrial TAs and the N-terminal small globular protein VHP (fig. S8B). We found that loss of MTCH2 by either knockout or knockdown affects the insertion of diverse mitochondrial TA proteins into the outer membrane, but not unrelated IMS- or matrix-targeted controls (Fig. 3A and fig. S9). Critically, reintroduction of MTCH2 restored insertion of mitochondrial TA substrates in our in vitro assay (Fig. 3B). We therefore concluded that MTCH2 is required for insertion of mitochondrial TAs into the outer membrane in a TMD-dependent manner.

We next tested whether MTCH2’s role in TA insertion involved a physical interaction with its substrates. To do this we adapted a site-specific crosslinking system using recombinant OMP25^BpA^ and mitochondria purified from human K562 cells (Fig. 3C; Table S3) (24). Crosslinked species were purified using an affinity tag on the N-terminus of OMP25^BpA^ and identified by mass spectrometry. In comparison to a non-UV activated control, we found that MTCH2 was highly enriched in our crosslinked samples. Consistent with these results, we could directly visualize a UV-dependent MTCH2-OMP25^BpA^ crosslinked species by western blotting (Fig. 3D). While TOM40 and CISD1 were enriched in our crosslinked samples, they were not significant hits in our genome-wide screen, and we have demonstrated TA insertion is a TOM-independent process (Fig. 1A and fig. S10). Taken together these data suggest that MTCH2 physically associates with nascent OMP25 during the insertion process.

Finally, to determine whether MTCH2 is sufficient for TA insertion, we purified MTCH2 from human cells (Fig. 3E) and optimized conditions for its reconstitution into liposomes (Fig. 3F). As a positive control, we also reconstituted the human EMC, which readily mis-inserts mitochondrial TAs (fig S11) (17), using a purification (25) and reconstitution strategy (26) as previously described. In a protease protection assay, OMP25 synthesized in native cytosol was inserted into MTCH2-containing proteoliposomes in a dose-dependent manner that correlated with MTCH2 concentration (Fig. 3F). OMP25 insertion was achieved at similar or greater efficiency than that observed with EMC proteoliposomes or isolated mitochondria. An ER TA, VAMP, showed no insertion across all conditions (fig. S11B). Because the protease protected fragment could be immunoprecipitated using a C-terminal affinity tag, and we observed no protected full-length product that would suggest translocation or leakiness of the liposomes, we concluded that MTCH2 inserts mitochondrial TAs in the correct orientation in this reconstituted system. Cumulatively, the requirement for MTCH2 in cells and in vitro for TA insertion, together with its reconstituted insertase activity and physical association with substrates, rigorously establish MTCH2 as an insertase for α-helical mitochondrial outer membrane proteins.

To reconcile these results with earlier observations that trypsin-treated mitochondria remain competent for TA insertion, we found that MTCH2 is largely trypsin resistant (Fig. 3G). We further verified that while trypsin treatment disrupted insertion of TOM-dependent substrates, insertion of OMP25 was minimally unaffected.

Bioinformatic analysis reveals that in addition to MTCH2, other examples of SLC25 family members lacking canonical sequence motifs are found in both mitochondria (MTCH1 and SLC25A46) and peroxisomes (PMP34) (Fig. 4A and fig. S12). Indeed, we found that depletion of the close paralog MTCH1 (27) by acute knockout (Fig. 4B and fig. S13A) or knockdown (fig. S13B) in a MTCH2 knockout background had an additive effect on biogenesis of many mitochondrial TAs. This result is consistent with recent pair-wise genetic interaction maps that have found a synthetic lethal relationship between MTCH1 and 2 (28), and the fact that MTCH1 was also a statistically significant hit in our genome-wide screen (fig. S12; S13C). We therefore propose that MTCH1 and MTCH2 are members of a unique class of membrane protein insertases that exploit the SLC25 transporter fold.

**Fig. 4.**
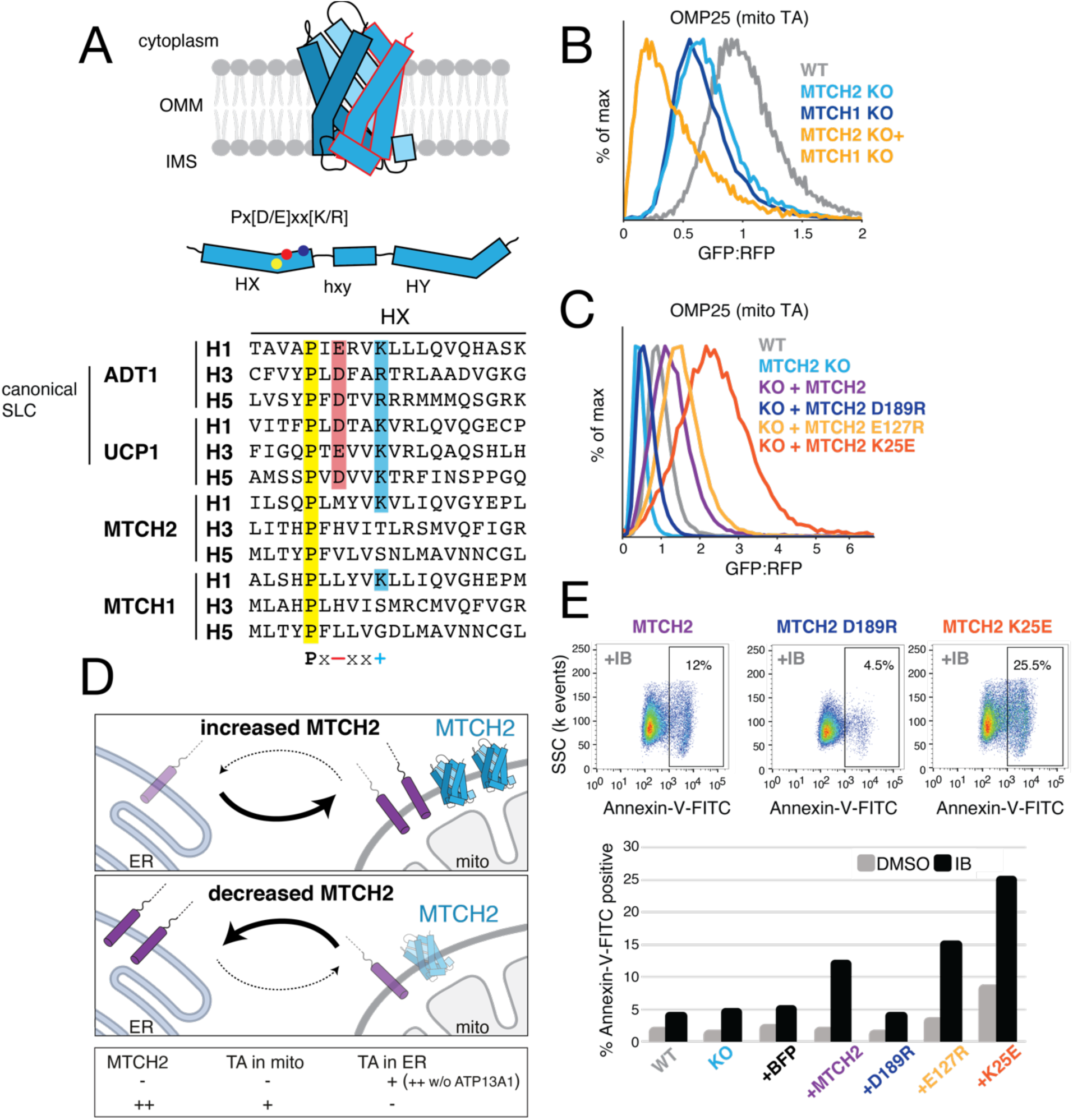
MTCH2 is a master regulator of outer membrane function. **(A)** Top: a predicted cartoon of MTCH2 based on the characteristic SLC25 TMD arrangement. SLC25 transporters are composed of three sets of two TMDs (six total), which are shown in unique shades of blue, with a single repeat outlined in red. In the middle, a schematic showing the location of the characteristic Px[D/E]xx[K/R] motif within a single, two helix, SLC25 repeat. Bottom: sequence alignment of all individual SLC25 repeats from two canonical inner membrane SLC25 transporters (ADT1, UCP1) and three diverged outer membrane SLC25 transporters (SLC25A46, MTCH1, MTCH2), with residues from the Px[D/E]xx[K/R] motif highlighted. **(B)** Flow cytometry analysis of OMP25-GFP11 integration into the outer membrane using the reporter assay described in Fig. 1B. To delineate the relationship between MTCH1 and 2 function in TA biogenesis, MTCH1 was depleted by transient knockout in either wild type (WT) or MTCH2 knock out (KO) cell lines. **(C)** As in (B) comparing cell lines with endogenously or exogenously overexpressed levels of MTCH2. The indicated mutations to MTCH2 that alter the electrostatic potential of its TMDs were tested for their effect on TA integration into the outer membrane. **(D)** Cell lines expressing GFP1-10 in the ER lumen were used to monitor mislocalization to the ER of mitochondrial TAs fused to a C-terminal GFP11. Table summarizing the analysis when either MTCH2 is depleted (in combination with the ER dislocase for mistargeted TAs, ATP13A1) or overexpressed (data in fig. S14, fig. S16, and fig. S17). **(E)** K562 cells expressing varying levels of MTCH2 or inactive (D189R) or hyperactive MTCH2 mutants (E127R or K25E; Fig. 4B) were treated with the chemotherapeutic imatinib mesylate (IB; 1 μM) or carrier (DMSO) for 72 hours. Apoptosis was assessed by staining with Annexin-V-FITC and analyzed by flow cytometry. Top: are representative dot plots displaying the fraction of apoptotic cells upon IB treatment while expressing WT MTCH2 compared to in inactive (D189R) or hyperactive mutant (K25E) Bottom: summary of apoptosis for all MTCH2 constructs in IB vs carrier treated control.

SLC25 transporters rely on charged and polar residues lining their central cavity to engage with substrates, suggesting MTCH2 may similarly depend on such interactions for TMD insertion (18). To test this, we identified a series of conserved charged and polar residues within MTCH2’s predicted TMDs and introduced mutations at positions that would therefore alter the electrostatic potential of its intramembrane surfaces. We found that while overexpression of wild type MTCH2 alone is sufficient to increase insertion of TA proteins into the outer membrane (fig. S14), a single charge-swap mutation strongly decreased its activity even when expressed at similar levels (Fig. 4C, fig. S15A and fig. S15B). Conversely, we also identified MTCH2 mutations that increase mitochondrial TA insertion, even when expressed ectopically in wild type cells that contain an endogenous copy of MTCH2 (Fig. 4C and fig. S15C). We therefore conclude that MTCH2’s role in TA insertion relies on the positioning of charged and polar residues within the bilayer, potentially an example of convergent evolution across many systems that move substrates into or across membranes (25, 29, 30). A significant number of mitochondrial proteins are enriched in basic residues immediately C terminal to their TMDs, and may be particularly reliant on charged surfaces along their route into the membrane (31).

Given MTCH2’s central role in mitochondrial TA biogenesis, we asked whether it may broadly affect cellular proteostasis. First, we reasoned that manipulating MTCH2 levels might affect the distribution of substrates between mitochondria and the ER, where they are known to mislocalize (32, 33). To test this, we adapted our split GFP system to report on TA integration into the ER by expressing GFP1-10 in the ER lumen (34). Indeed, depletion of MTCH2 led to an increase in ER insertion for a variety of mitochondrial TAs both in cells and in vitro (fig. S16A and fig. S16B). This effect was enhanced by further depleting ATP13A1, a recently described quality control factor that dislocates mislocalized mitochondrial TA proteins from the ER membrane for degradation by the ubiquitin proteasome pathway (24) (fig. S16C). Conversely, increased expression of MTCH2 led to less OMP25-GFP11 in the ER (fig. S17). The insertion of ER resident TA proteins was unaffected by MTCH2 levels, suggesting it acts as part of an orthogonal pathway specifically catering to mitochondrial TA biogenesis (fig. S16A and fig. S17B). These data suggest that MTCH2 is the central ‘gatekeeper’ for the outer membrane: MTCH2 levels and activity dictate the cytosolic reservoir of mitochondrial TAs, which then can be re-routed to the ER if unable to successful integrate into mitochondria (Fig. 4D).

Finally, considering that insertion of several MTCH2-dependent TAs plays a central role in cellular apoptosis, we reasoned that MTCH2 activity may affect cellular sensitivity to apoptotic stimuli. To test this, we overexpressed MTCH2 (along with a BFP marker) or a BFP control in human K562 cells, which are derived from a myelogenous leukemia cell line known to upregulate the anti-apoptotic TA, BCL2L1 (35). We treated cells with imatinib, a tyrosine-kinase inhibitor commonly used as a targeted treatment for several cancers, including leukemias, and measured apoptosis by surface staining with Annexin V. Because MTCH2 substrates include both the anti- and pro-apoptotic TAs (i.e. BCL2L1 and BAK1; Fig. 3A), we did not a priori know how modulation of MTCH2 activity would affect apoptosis propensity, and indeed may be cell-type and environment specific. However, we found that while knockout of MTCH2 did not appreciably alter apoptosis propensity in this system (Fig. 4E), overexpression of wild type MTCH2 markedly sensitizes K562 cells to imatinib treatment, leading to a ∼2-fold increase in apoptotic cells compared to a BFP control. Critically, this sensitization appears to depend on MTCH2’s insertase activity: expression of an inactive mutant had no effect, while the hyperactive mutant (K25E) enhanced K562 apoptosis propensity both with and without drug treatment. These results are consistent with MTCH2’s central role in insertion of pro-apoptotic mitochondrial TAs into the outer membrane, a definitive early step in the apoptosis pathway.

In summary, we have demonstrated that MTCH2 is a defining member of a family of membrane protein insertases that are essential for viability, and necessary and sufficient for insertion of TAs into human mitochondria. MTCH2’s insertase activity relies on hydrophilic residues within its TMDs, a common feature of many membrane protein translocases including the EMC (25, 30, 36), Hrd1 (37), and YidC (38). MTCH2’s role also appears to extend to the integration of a broader class of α-helical proteins into the outer membrane, including signal anchored and multipass proteins, which remains to be systematically delineated. Homologs of MTCH2 are present in metazoan peroxisomes and its orthologs are found throughout holozoa, suggesting that the MTCH2 family has co-opted the SLC25 transporter fold to function in diverse biological membranes. This type of functional divergence has been previously described for SFXN4 and ATP13A1, both transporter family proteins that have evolved biogenesis or dislocase functions, respectively (18, 24, 39).

We have shown that MTCH2 is involved in modulating fundamental cellular phenomena including protein localization and apoptosis. Previously, loss of MTCH2 has been reported to lead to a diverse range of phenotypes including dysregulation of mitophagy, mitochondrial fragmentation, recruitment of tBID, and altered lipid homeostasis, and was also identified in a recent genome-wide association study for Alzheimer’s disease (40–42). The identification of MTCH2 as a key gatekeeper for α-helical outer membrane proteins now provides a molecular explanation for its pleotropic phenotypes, many of which can be directly ascribed to defects in biogenesis of MTCH2 substrates. Identifying the molecular role of MTCH2 raises the potential for manipulating its function to affect proteostasis and cell fate, especially in the context of disease.

## Acknowledgements

We thank T. Pleiner and Z. Levine for careful reading and input on the manuscript. We thank: the Whitehead Institute Flow Cytometry Core and Kathy Daniels for access to FACS machines; the Whitehead Institute Genome Technology Core for support with sequencing of screen libraries; the Caltech Flow cytometry facility; and the Ting-Yu Wang and the Proteome Exploration Laboratory at Caltech for support for mass spectrometry.

## Materials and Methods

### Plasmids and antibodies

Endogenous sequences used in this study for in vitro and in vivo analysis were sourced from UniProtKB/Swiss-Prot and included: squalene synthase isoform 1 (SQS/FDFT1; Q6IAX1), synaptojanin-2 binding protein (OMP25/SYNJBP; P57105-1), mitochondrial antiviral-signaling protein (MAVS; **Q7Z434-1**), mitochondrial import inner membrane translocase subunit Tim9 (TIM9; Q9Y5J7-1), vesicle associated membrane protein 2 (VAMP; **P51809-1**), FUN14 domain-containing protein 1 (FUNDC1; Q8IVP5-1), mitochondrial import receptor subunit TOM5 homolog (TOM5; **Q8N4H5-1**), mitochondrial import receptor subunit TOM22 homolog (TOM22; Q9NS69-1), ubiquitin carboxyl-terminal hydrolase 30 (USP30; Q70CQ3-1), apoptosis regulator BAX (BAX; **Q07812-1**), Bcl-2 homologous antagonist/killer (BAK1; **Q16611-1**), Bcl-2-like protein 1 (BCL2L1; **Q07817-1**), Cytochrome b5 type B (CYB5B; O43169-1), peptidyl-prolyl cis-trans isomerase FKBP8 (FKBP8; **Q14318-1),** mitochondrial fission factor (MFF; **Q9GZY8-1),** inactive hydroxysteroid dehydrogenase-like protein 1 (HSDL1; **Q3SXM5-1**), amine oxidase [flavin-containing] A (MAOA; **P21397-1**), amine oxidase [flavin-containing] B (MAOB; **P27338-1**), mitochondrial Rho GTPase 1 (RHOT1; **Q8IXI2-1**), mitochondrial Rho GTPase 2 (RHOT2; **Q8IXI1-1**), and mitochondrial carrier homolog 2 (MTCH2; Q9Y6C9-1).

Constructs for expression in rabbit reticulocyte lysate (RRL) used the SP64 vector as a backbone (Promega). For in vitro insertion reactions, an N-terminal 3xFLAG tag and a C-terminal 6xHis tag were appended to the respective terminus for affinity purification (see fig. S8A). Where noted, the transmembrane domain (TMD) of TA proteins along with N- and C-terminal flanking sequences were instead conjugated to maltose binding protein (MBP) (45) or villin headpiece (VHP) domains as illustrated in fig. S8B. When monitoring insertion into the endoplasmic reticulum (fig. S4 and S16), constructs were modified to replace the 6xHis tag with a C-terminal opsin tag containing a glycosylation acceptor site which can be used as a proxy for insertion (46). TOM dependent mitochondrial import substrates were designed by fusing dihydrofolate reductase (DHFR) to mitochondrial transit sequences from either *N. crassa* ATP synthase subunit 9 (residues 1-69) or *S. cerevisiae* cytochrome b_2_ (residue 1-167) which direct import to the mitochondrial matrix or inner membrane space (IMS), respectively.

For expression in K562 cells, the basis for all constructs was a mammalian expression lenti-viral backbone containing a UCOE-EF-1α promoter and a 3’ WPRE element ((47); Addgene #135448). The exception was the dual fluorescent reporter used for the CRISPRi screen (RFP-P2A-OMP25-GFP11) which was integrated into an SFFV-tet3G backbone (48). The dual color reporter system used for in vivo experiments has been previously described (49) (26). Note that the mCherry variant of RFP was used in all instances, but the simpler nomenclature of RFP is used in the text and figures. For complementation with the GFP1-10 system, the GFP11 tag (RDHMVLHEYVNAAGIT) was appended to the appropriate terminus of the indicated protein as determined by predicted topology (see Fig. 2C). In order to express GFP1-10 in the ER lumen, the human calreticulin signal sequence was appended preceding GFP1-10-KDEL (50) (51). For targeting to the inner membrane space of the mitochondria, either the targeting signal from MICU1 (aa.1-60) (14) or the targeting sequence from LACTB (aa 1-68) (44) was appended to the N terminus of GFP1-10. Expression of MTCH2 or indicated mutants was from a BFP-P2A-(MTCH2) cassette, allowing us to gate and sort for expressing cells.

The single sgRNA for MTCH2 (GACGGAGCCACCAAGCGACC) was generated by annealed oligo cloning of top and bottom oligonucleotides (Integrated DNA Technologies, Coralville, IA) into a lentiviral pU6-sgRNA EF-1α-Puro-T2A-BFP vector digested with BstXI/BlpI (Addgene, cat# 84832). BFP was excised in certain sgRNA variants when the color interfered. Though most experiments were done with a single guide that gave robust knock-down, key results were verified with an additional guide (GGGCTCACCGGGTCGCTTGG) to exclude possible off target effects. In many instances, we used a programmed dual sgRNA guide vector ((52); Addgene #140096) to allow for multiple genes to be depleted at once or to increase efficiency of knock-down. Dual guide pairs included MTCH2-ATP13A1 (GACGGAGCCACCAAGCGACC, GGGTAAAGCAGCCCGGCGAA), MTCH1-MTCH1 (GCGGCACCGCCGCGAGCCCA, GAGCCCAGGGCGCCACTTCC), and MTCH2-EMC2 (GACGGAGCCACCAAGCGACC, GGAGTACGCGTCCGGGCCAA). Transient knock-out of MTCH1 was achieved by modifying a pLentiCRISPR backbone ((53), Addgene #102315) to express the following guides: GGACAACGCCCCGACCACTG and CTGCATCATCATCTCGTAGG.

Constructs for recombinant bacterial protein expression were all cloned in the pQE plasmid (Qiagen). Su9-DHFR and CaM-3C-Alfa-Sec61β(2–60)-OMP25(112–145)F128Amber were cloned downstream of a His_14_-*bd*SUMO tag. A GFP Nb fusion protein used in this manuscript was modified from a previously established construct ((25); Addgene ID #149336) to exclude the SUMO^Eu1^ tag upstream of the GFP Nb. Constructs for the expression of SENP^EuB^ ((54); Addgene ID #149333), *bd*SENP1 ((55); Gift from Dirk Görlich; Addgene ID #104962), and BirA ((56); Gift from Dirk Görlich, Addgene ID #149334), and for site-specific incorporation of BpA at the Amber codon position ((57); Addgene #31190) have all been previously described.

Constructs for generating human stable cell lines for recombinant protein expression used either the pHAGE2 plasmid (Gift from Magnus A. Hoffmann and Pamela Bjorkman) for lentiviral integration into the Expi293F cell line or the pcDNA5/FRT/TO plasmid (Thermo Scientific Cat. #V652020) for recombinase-mediated integration into Flp-In 293 T-REx cell line. MTCH2 was N-terminally fused with a GFP-SUMO^Eu1^ tag and downstream of a doxycycline-inducible CMV promoter. The EMC3-GFP-P2A-RFP expression vector was previously described ((25)).

The following antibodies were used in this study: MTCH2 (ab113707, Abcam, UK); FUNDC1 (OAAB12808, Aviva systems biology, USA); CYB5B (HPA007893, Atlas antibodies, USA); MIRO2 (RHOT2) (ab224089, Abcam, UK); CYC1 (4272, Cell signaling technology, USA); SYNJ2BP (OMP25) (15666-1-AP, Proteintech, USA); EMC3 (67205, Proteintech, USA); VDAC1 (sc-390996, Santa Cruz Biotech, USA); mitofilin (ab110329, Abcam, UK); SAMM50 (ab133709, Abcam, UK); ATP13A1 (16244-1-AP, Proteintech, USA); tubulin (T9026, Sigma-Aldrich, USA). The ALFA tag was detected by coupling HRP to an ALFA nanobody (58). The Sec61β antibody was a gift from Ramanujan Hegde. Secondary antibodies used for Western blotting were: Goat anti-mouse- and anti-rabbit-HRP (#172-1011 and #170-6515, Bio-Rad, USA).

### Cell culture and cell line generation

K562 cells were grown in RPMI-1640 with 25 mM HEPES, 2.0 g/L NaHCO_3_, and 0.3 g/L L-glutamine supplemented with 10% FBS (or Tet System Approved FBS), 2 mM glutamine, 100 units/mL penicillin, and 100 μg/mL streptomycin. Cells were maintained at a confluency between 0.25-1 x10^6^ cells/mL. HEK293T cells were grown in DMEM supplemented with 10% FBS, 100 units/ml penicillin and 100 μg/ml streptomycin. All cell lines were grown at 37°C.

For the expression of EMC3-GFP, Flp-In 293 T-Rex cells were purchased from ThermoFisher Scientific (USA) (RRID: CVCL_U427) and grown in DMEM supplemented with 2 mM glutamine, 10% FBS, 15 µg/mL Blasticidine S, and 100 µg/mL Zeocin. The open reading frame to be integrated into the genomic FRT site was cloned into the pcDNA5/FRT/TO vector backbone and cell lines were generated according to the manufacturer’s protocol. To allow for large scale growth, these cells were adapted to grow in suspension. Briefly, over the course of 10 days the FBS-supplemented DMEM was serially diluted with FreeStyle 293 Expression Medium (ThermoFisher Scientific, USA). Once growing in 100% FreeStyle Medium, the cells were transferred to 1-2 L roller bottles (Celltreat, USA) and grown in a shaking incubator operating at 8% CO_2_ and rotating at 125 rpm. For the expression of MTCH2, lenti-viral infected inducible Expi293F suspension cells were grown in Expi293 expression media (ThermoFisher Scientific, USA) at 37°C, 8% CO_2_ and 125 rpm shaking in 1 L roller bottles with vented caps (Celltreat, USA)

Three K562 cell lines were used a basis for cell lines generated in this study: CRISPRi K562 cells expressing dCas9-BFP-KRAB (KOX1-derived) (15), K562-dCas9-BFP-KRAB Tet-On cells (48), and CRISPRi cells generated by stably expressing ZIM3 KRAB-dCas9-P2A-BFP from a UCOE-SFFV promoter (22). To generate cell lines with GFP1-10 in the mitochondrial IMS or ER lumen, virus was made from the respective constructs: LACTB(GFP1-10), MICU1(GFP1-10) or CalR(GFP1-10)-KDEL. CRISPRi K562 cells were infected with lenti-virus and sorted into 96-well plates as single cell clones using a Sony Cell Sorter (SH800S). After expansion, correct cell lines were confirmed by successful complementation with a construct targeted to the respective compartment, appended to a GFP11. To generate the cell line used for screening, lenti-virus containing MICU1(GFP1-10) and RFP-P2A-OMP25-GFP11 under a tet-inducible promoter were co-infected to one copy per cell in CRISPRi K562 Tet-On cells, single cell sorted, and verified by induction with doxycycline (100 ng/uL) and microscopy in conjunction with MitoTracker staining to confirm correct localization.

To generate MTCH2 knock-out cell lines, K562 CRISPRi cells with or without MICU1(GFP1-10) were nucleofected with a MTCH2 targeting guide in the pX458 backbone (Addgene plasmid # 48138) using the Lonza SF Cell Line 96-well Nucleofector Kit (V4SC-2096). The pX458 backbone was adapted to express two sgRNAs targeting MTCH2 [AGCCGACATGTCTCTAGTGG], [GGCTTTGCGAGTCTGAACGT]. Two days following nucleofection, GFP-positive cells were single cell sorted into 96-well plates. After colonies from single cells grew out, loss of MTCH2 was confirmed by Western blotting. CRISPR-Cas9-induced genome edits were identified using the computational pipeline described in (59).

### Lentivirus

Lentivirus was generated by co-transfecting HEK293T cells with two packaging plasmids (pCMV-VSV-G and delta8.9, Addgene #8454) and the desired transfer plasmid using TransIT-293 transfection reagent (Mirus). 48 hours after transfection, the supernatant was collected and flash frozen. In all instances, virus was rapidly thawed prior to transfection. Virus for the genome-wide CRISPRi screen was also generated using this method.

### CRISPRi KD Screen

The genome-scale CRISPRi screen was performed in duplicate as previously described (15) (60). The hCRISPRi-v2 compact library (5 sgRNAs per gene, Addgene pooled library #83969) was transduced in duplicate into 330 million K562-CRISPRi-Tet-ON-((MICU1)-GFP1-10)-(tet-RFP-P2A-OMP25-GFP11) cells at multiplicity of infection (MOI) < 1 (percentage of transduced cells 48 hours after infection as measured by BFP positive cells: 30-35%). Cells were grown in 1 L of media in 1 L spinner flasks (Bellco, SKU: 1965-61010). 48 hours after spinfection with the genome-wide library, guide positive cells were selected with 1 µg/mL puromycin for three days. Following a 36 hour recovery, cells were induced with 100 ng/mL doxycycline for 36 hours and sorted on a FACS AriaII Fusion Cell Sorter. To ensure that the culture was maintained at an average coverage of more than 1000 cells per sgRNA, cells were diluted daily to 0.5x10^6^ cells/mL.

During sorting, cells were gated for BFP (indicating a guide-positive cell), as well as GFP and RFP signal (successfully induced). Cells were sorted based on the GFP:RFP ratio of this final gated population. Roughly 40 million cells with either the highest (30%) or lowest (30%) RFP:GFP ratio were collected, pelleted and flash-frozen. Genomic DNA was purified using the Nucleospin Blood XL kit (Takara Bio, #740950.10) and amplified by index PCR with barcoded primers. The resulting guide library (∼264 bp) was purified using SPRIbeads (SPRIselect Beckman Coulter #B23318). Sequencing was performed using an Illumina HiSeq2500 high throughput sequencer. Sequencing reads were aligned to the CRISPRi v2 library sequences, counted and quantified (60). Generation of negative control genes and calculation of phenotype scores and Mann-Whitney p-values was performed as described previously (15) (60). Gene-level phenotypes and counts are available in Supplementary Table 1.

### Protein expression and purification

#### Su9-DHFR and CaM-3C-Alfa-Sec61β-OMP25(BpA)

The BL21(DE3) expression strain was used to express Su9-DHFR and CaM-3C-Alfa-Sec61β-OMP25(BpA) in LB media. Su9-DHFR expressing cultures were induced with 1 mM IPTG after an optical density of 0.6 was reached. After induction, Su9-DHFR was expressed at 37°C for 3 hours. CaM-3C-Alfa-Sec61β-OMP25(BpA) was co-transformed with pEVOL-BpF and grown to an optical density of 0.2 followed by induction with 1% arabinose and the addition of 1 mM BpA (Bachem). Cells were then grown to an optical density of 0.6 followed by induction with 1 mM IPTG and expression at 25°C for 6 hours. Cells were pelleted by centrifugation and resuspended in a lysis buffer containing 500 mM NaCl, 50 mM Tris pH 7.5, 10 mM imidazole, 5 mM β-ME.

For purification, *E. coli* resuspensions were supplement with EDTA-free protease inhibitor tablets (Roche) and lysozyme prior to lysis via sonication. Lysate was clarified by centrifugation at 18,000 rpm for 30 minutes in an SS-34 rotor. Clarified lysate was incubated with NiNTA resin for 30 minutes while rolling at 4°C. NiNTA resin was washed extensively with resuspension buffer, and then equilibrated with a SENP elution buffer containing 150 mM NaCl, 50 mM Tris pH 7.5, 10 mM imidazole, 5 mM β-ME, 10% glycerol. NiNTA resin was then incubated with *bd*SENP1 for 2 hours at 4°C to release SUMO-cleaved protein from the resin.

For CaM-3C-Alfa-Sec61β-OMP25(BpA) the resuspension buffer included 1 mM CaCl_2_, and SENP elution buffer contained 100 mM NaCl, 50 mM Tris pH 7.5, 10 mM imidazole, 5 mM β-ME, 1 mM CaCl_2_. Additionally, protein was cleaved overnight with 3C protease following SENP1 elution. Cleaved protein was then concentrated to 250 µL and injected onto a Superdex 200 increase 10/300 GL size exclusion column equilibrated in a buffer containing 150 mM KoAc, 50 mM HEPES pH 7.4, 2 mM MgoAc_2_, 1 mM CaCl_2_, 1 mM DTT, 10% glycerol. Protein-containing fractions were pooled, concentrated, aliquoted and flash frozen.

### Biotinylated GFP-Nb and Alfa-NB

Expression and purification of all GFP and Alfa nanobody constructs as well as *bd*SENP1, BirA, and SENP^EuB^ generally proceeds as follows: the NEB Express I^q^ expression strain was used with TB medium. Cultures were induced with 0.2 mM IPTG after an optical density of 2.0 was reached. Protein was expressed at 18°C for 18-20 hours. Cells were pelleted by centrifugation and resuspended in a lysis buffer containing 50 mM Tris pH 7.5, 300 mM NaCl, 20 mM imidazole, 1 mM DTT, 1 mM PMSF. Cells were lysed by sonication and lysate was clarified by centrifugation at 18,000 rpm for 30 minutes in an SS-34 rotor. Clarified lysate was incubated with NiNTA resin for 1 hour while rolling at 4°C. NiNTA resin was washed extensively with resuspension buffer and the eluted with a buffer containing 50 mM Tris pH 7.5, 300 mM NaCl, 500 mM imidazole, 250 mM sucrose. The imidazole was removed using a PD-10 desalting column (GE Healthcare, USA). The following protein-specific modifications were applied to the above protocol: the Rosetta-gami 2(DE3) expression strain was used to express His_14_-Avi-SUMOstar-AlfaNb and expression time was limited to 6 hours at 18°C. SENP^EuB^ was limited to an expression time of 6 hours at 18°C. *bd*SENP1 was expressed as a His_14_-TEV-fusion and was cleaved with TEV protease overnight after buffer exchange and then ran over NiNTA resin to remove the cleaved tag. The expression and purification of His_14_-Avi-SUMO^Eu1^-anti GFP nanobody, *bd*SENP1, BirA, and SENP^EuB^ have all (61) been previously described (25) (54).

All nanobody constructs used in this publication were biotinylated in a buffer containing 50 mM Tris pH 7.5, 100 mM NaCl, 12.5 mM MgCl_2_, 10 mM ATP, 10 mM biotin and BirA at a 1:50 molar ratio to the nanobody substrate. Biotinylation reactions were incubated at 25°C for 3 hours and then buffer exchanged in a PD-10 column equilibrated with a buffer containing 50 mM Tris pH 7.5, 200 mM NaCl, 1 mM DTT, 250 mM sucrose. Biotinylated protein was aliquoted and flash frozen.

### MTCH2 and EMC

We isolated MTCH2 and EMC (via EMC3-GFP) under native conditions from detergent-solubilized cells using a biotinylated anti-GFP nanobody, expressed and purified as previously described (25). Cells were grown in 1 L roller bottles. For the MTCH2 lines, expression was induced for at least 48 hours with 1 μg/mL doxycycline. Cells were then harvested by centrifugation, washed with 1x PBS, and then pellets were weighed.

Purification generally proceeded as follows: cell pellets were resuspended at a ratio of 1 g to 10 mL hypotonic lysis buffer containing 10 mM HEPES pH 7.5, 10 mM KoAc, 0.15 mM MgoAc_2_, 0.5 mM DTT, supplemented with EDTA-free protease inhibitor tablets (Roche). The cell resuspension was incubated on ice for 10 minutes to allow cells to swell, and then lysed in a Dounce homogenizer with 10x strokes. The NaCl concentration was adjusted to 180 mM immediately after Dounce homogenization. Cell membranes were pelleted by centrifugation at 18,000 rcf in an SS-34 (28020TS, Thermo Fisher) rotor for 10 minutes. Supernatant was discarded and cell membranes were washed by resuspending and pelleting 2x in membrane wash buffer containing 10 mM HEPES/KOH pH 7.5, 200 mM NaCl, 0.15 mM MgoAc_2_, 0.5 mM DTT. The resulting pellet was resuspended at a ratio of 1 g (original cell pellet weight) to 6.8 mL solubilization buffer containing 50 mM HEPES pH 7.5, 200 mM NaCl, 2 mM MgoAc_2_, 1% deoxy-BigCHAP (DBC; Anatrace Cat # B310), 1 mM DTT, supplemented with EDTA-free protease inhibitor tablets (Roche). After 30 minutes of head-over-tail incubation with solubilization buffer, the lysate was cleared by centrifugation for 30 minutes at 4°C and 18,000 rpm in a SS-34 rotor. The supernatant was then added to pre-equilibrated magnetic Streptavidin resin (ThermoFisher) bound to biotinylated anti-GFP nanobody and blocked with free biotin. After 1 hour of binding while rolling at 4°C, the resin was washed four times with wash buffer (solubilization buffer with 0.2% DBC). Resin was then incubated in wash buffer + 600 nM SENP^EuB^ on ice for 2 hours to release SUMO-cleaved MTCH2 from the resin. Eluted samples were analyzed via SDS-PAGE with Sypro Ruby stain (Bio-Rad).

For EMC3-GFP, the whole cell pellet was solubilized in 1% DBC-containing buffer, without a hypotonic lysis and membrane washing step, and the GFP Nb was fused to a cleavable SUMO^Eu1^ module.

### Mitochondrial isolation and semi-permeabilized cells

Mitochondrial isolation for K562 cells was adapted from an established protocol (62). K562 cells were centrifuged at 220*g* for 5 minutes. Pellets were washed once in PBS and pelleted by spinning at 500*g* for 5 minutes. Pellets were resuspended in a homogenization buffer containing 210 mM mannitol, 70 mM sucrose, 5 mM HEPES pH 7.4, 10 mM EDTA, 1 mM PMSF, and 2 mg/mL BSA. After incubating on ice for 10 minutes, cells were then lysed with a glass Dounce homogenizer with a tight-fitting pestle, or a Potter-Elvehjem homogenizer motor-driven at 1600 rpm for large scale purifications. Homogenized cells were pelleted at 1300*g* for 5 minutes to remove nuclei and unbroken cells, then the supernatant was transferred to a clean tube. This step was repeated twice. Nuclei-free homogenized cells were then centrifuged at 11,000*g* for 10 minutes. The supernatant was removed and the mitochondria-containing pellet was then resuspended in an isolation buffer containing 210 mM D-mannitol, 70 mM sucrose, 5 mM HEPES pH 7.4, 10 mM EDTA. Mitochondria were then pelleted and resuspended in fresh isolation buffer to wash away BSA and cytoplasmic proteins. After a final pelleting step, mitochondria were resuspended in a small volume (5-50 µL) of isolation buffer. To normalize mitochondrial samples, the protein concentration was measured using a Bradford assay.

For experiments using trypsin-treated mitochondria, the final 2 was steps performed after pelleting mitochondria used import buffer containing 250 mM sucrose, 5 mM MgoAc2, 80 mM KoAc, and 20 mM HEPES. Mitochondria were then mixed with 0 or 50 µg/ml trypsin (Sigma-Aldrich Cat# T1426) dissolved in import buffer and incubated on ice for 30 minutes. 50 µg/ml trypsin inhibitor (Sigma-Aldrich Cat# T9128) and 1 mM PMSF were added to quench the reaction. Mitochondria were pelleted, washed once in import buffer + 5 µg/ml trypsin inhibitor, and then resuspended in a concentrated volume prior to use in import experiments.

To further enrich mitochondrial samples for certain experiments, a percoll gradient was used. Isolation buffer density gradients were formed in 3 layers with 40%, 26%, and 12% percoll. Resuspended mitochondria were layered on top of the gradient. Gradients were centrifuged at 45,000*g* for 45 minutes in a TLS-55 rotor (Beckman Coulter). Pure mitochondria were retrieved from the 40%-26% percoll interface (see fig. S1B). Mitochondria were diluted 5-fold in isolation buffer, and then pelleted and washed in isolation buffer twice more.

HEK293T cells, either wild-type or an EMC5 knock-out background (26) were semi-permeabilized using standard methods (16). Briefly, 3x10^6^ cells were collected and washed once with ice cold wash buffer containing 25 mM HEPES pH 7.4, 100 mM KoAc, 2 mM MgoAc2. The cells were resuspended in 1 mL SP buffer containing 25 mM Hepes pH 7.4, 100 mM KoAc, 2 mM MgoAc2, 50 µg/mL digitonin) and incubated on ice for 5 minutes. The semi-permeabilized cells were collected by centrifugation at 500*g* for 5 minutes at 4°C, with the digitonin removed by washing three times. Finally, the cells were pelleted at 12,000*g* for 15 seconds and resuspended in 10 µL wash buffer. To test the integrity of the outer mitochondria membrane as in fig. S1A, SP cells and purified mitochondria (as described above) were incubated with the indicated amount of proteinase K (PK). Following quenching, the resulting reaction was subjected to blotting against mitofilin, an inner membrane localized protein which should not be accessible by PK when the outer mitochondrial membrane is intact.

### In vitro translation and insertion

In vitro translations were carried out in rabbit reticulocyte lysate (RRL) as previously described (26). Constructs for in vitro translation reactions were based on the SP64 vector (Promega, USA). Templates for transcription were generated by PCR, with primers binding and upstream of the SP64 promoter and roughly 200 bp downstream of the stop codon (63). Following transcription at 37°C for 1.5 hours, reactions were used directly in a translation reaction. Substrates were translated for 15-30 minutes at 32°C in the presence of radioactive ^35^S-methionine. Prior to addition of mitochondria or semi-permeabilized cells, 1 mM puromycin was added to prevent further synthesis.

Mitochondrial insertion reactions used isolated mitochondria, prepared as described above. Insertion reactions were performed by diluting 4 μL of a puromycin treated translation reaction in 50 μL of import buffer (250 mM sucrose, 5 mM MgoAc2, 80 mM KoAc, 20 mM HEPES pH 7.4, 2.5 mM ATP, 15 mM succinate) with 15 μg of purified mitochondria and further incubating at 32°C for 30 minutes. For competition experiments in Fig. 1A, insertion reactions were carried out in the presence of 1, 2, or 5 μM recombinant Su9-DHFR, and 5 μM methotrexate (BP266510, Fisher Chemicals, USA).

Protease digestions were initiated by the addition of proteinase K at 0.25 mg/mL, and reactions were then incubated on ice for 1 hour. Reactions were quenched by the addition of 5 mM PMSF in DMSO, followed by transfer to boiling 1% SDS (final concentration) in 0.1 M Tris/HCl pH 8.0. His-tagged protected fragments were enriched by incubating with NiNTA resin in IP buffer (50 mM HEPES pH 7.5, 500 mM NaCl, 10 mM imidazole, 1% Triton). Proteinase K digested reactions were diluted to 1 mL and mixed with 10 μL resin, then incubated with end-over-end mixing for 1.5 hours at 4°C. The resin was further washed with 3x1 mL IP buffer, and the products eluted from the resin in sample buffer containing 50 mM EDTA pH 8.0.

Insertion reactions with semi-permeabilized (SP) cells used a ratio of 1 μL cells per 10 μL translation reaction. To verify insertion of substrates into the ER as indicated by glycosylation, a tripeptide competitor of glycosylation (Asn-Tyr-Thr) was added at 50 μM when indicated.

### Mass spectrometry

For the crosslinking-IP samples, TCA-precipitated pellets were resuspended in a buffer containing 8 M Urea and 100 mM Tris pH 8.5. The sample was reduced by incubation with 3 mM TCEP for 20 minutes, then alkylated by incubation with 10 mM iodoacetamide for 15 minutes, all at room temperature. The sample was then digested with 2 ng/µL LysC for 4 hours at room temperature, diluted 4-fold with 100 mM Tris pH 8.5 and CaCl_2_ was added to 1 mM. The sample was then digested with 4 ng/µL Trypsin overnight at room temperature. Samples were acidified by adding trifluoroacetic acid to 0.5%, desalted using Pierce C18 spin columns (Pierce), lyophilized, and then resuspended in 2% acetonitrile, 0.2% formic acid.

For the mitochondrial proteomics experiments, the S-trap sample preparation kit (ProtiFi) was used according to the manufacturer’s instructions. Sample was digested on the S-trap column with 1 µg Trypsin per 10 µg protein overnight at 37°C. In addition to the provided S-trap sample preparation protocol, a final elution step with 70% acetonitrile, 1% formic acid was added.

Eluted peptides were lyophilized, and then resuspended in 2% acetonitrile, 0.2% formic acid. The peptide concentration was determined with a Quantitative Fluorometric Peptide Assay (Pierce) kit.

LC-MS/MS analysis for the crosslinking-IP experiment was performed with an EASY-nLC 1200 (ThermoFisher Scientific, San Jose, CA) coupled to a Q Exactive HF hybrid quadrupole-Orbitrap mass spectrometer (ThermoFisher Scientific, San Jose, CA). Peptides were separated on an Aurora UHPLC Column (25 cm × 75 μm, 1.6 μm C18, AUR2-25075C18A, Ion Opticks) with a flow rate of 0.35 μL/min for a total duration of 75 min and ionized at 1.8 kV in the positive ion mode. The gradient was composed of 2-6% solvent B (3.5 min), 6-25% B (42 min), 25-40% B (14.5 min), and 40–98% B (15 min); solvent A: 2% ACN and 0.2% formic acid in water; solvent B: 80% ACN and 0.2% formic acid. MS1 scans were acquired at the resolution of 60,000 from 375 to 1500 m/z, AGC target 3e6, and maximum injection time 15 ms. The 12 most abundant ions were targeted for MS2 scans acquired at a resolution of 30,000, AGC target 1e5, maximum injection time of 60 ms, and normalized collision energy of 28. Dynamic exclusion was set to 30 s and ions with charge +1, +7, +8 and >+8 were excluded. The temperature of ion transfer tube was 275°C and the S-lens RF level was set to 60. MS2 fragmentation spectra were searched with SEQUEST running within Proteome Discoverer (version 2.5, Thermo Scientific) against the UniProt human reference proteome comprised of 79,052 proteins covering 20,577 genes (UP000005640). The maximum missed cleavages was set to 2. Dynamic modifications were set to oxidation on methionine (M, +15.995 Da), deamidation on asparagine and glutamine (N and Q, +0.984 Da), phosphorylation of serine and threonine (S and T, +79.966 Da), protein N-terminal acetylation (+42.011 Da), and protein N-terminal Met-loss (-131.040 Da). Carbamidomethylation on cysteine residues (C, +57.021 Da) was set as a fixed modification. The maximum parental mass error was set to 10 ppm, and the MS2 mass tolerance was set to 0.03 Da. The false discovery threshold was set strictly to 0.01 using the Percolator Node validated by q-value. The relative abundance of parental peptides was calculated by integration of the area under the curve of the MS1 peaks using the Minora LFQ node.

To identify enriched proteins, proteins that were detected in both samples were ranked by iBAQ intensity within each sample, and enrichment was assessed based on the difference between the +UV and the -UV iBAQ rank. The final results of this analysis are listed in Supplementary Data 3.

LC-MS/MS analysis to assess differences in protein content between percoll gradient-enriched mitochondria derived from K562 wild-type or MTCH2 depleted cells was performed with an EASY-nLC 1200 (ThermoFisher Scientific, San Jose, CA) coupled to an Orbitrap Eclipse Tribrid mass spectrometer (ThermoFisher Scientific, San Jose, CA). Peptides were separated on an Aurora UHPLC Column (25 cm × 75 μm, 1.6 μm C18, AUR2-25075C18A, Ion Opticks) with a flow rate of 0.35 μL/min for a total duration of 75 min and ionized at 1.8 kV in the positive ion mode. The gradient was composed of 2-6% solvent B (3.5 min), 6-25% B (42 min), 25-40% B (14.5 min), and 40–98% B (15 min); solvent A: 2% ACN and 0.2% formic acid in water; solvent B: 80% ACN and 0.2% formic acid. MS1 scans were acquired at the resolution of 120,000 from 350 to 1,600 m/z, AGC target 1e6, and maximum injection time of 50 ms. MS2 scans were acquired in the ion trap using fast scan rate on precursors with 2-7 charge states and quadrupole isolation mode (isolation window: 0.7 m/z) with higher-energy collisional dissociation (HCD, 30%) activation type. Dynamic exclusion was set to 30 s. The temperature of ion transfer tube was 300°C and the S-lens RF level was set to 30. MS2 fragmentation spectra were searched with SEQUEST running within Proteome Discoverer (version 2.5, Thermo Scientific) against the reviewed sequences from the UniProt human reference proteome comprised of 20,361 proteins (UP000005640). The maximum missed cleavages were set to 2. Dynamic modifications were set to oxidation on methionine (M, +15.995 Da), deamidation (N and Q, +0.984 Da), protein N-terminal acetylation (+42.011 Da) and protein N-terminal Met-loss (-131.040 Da). Carbamidomethylation on cysteine residues (C, +57.021 Da) was set as a fixed modification. The maximum parental mass error was set to 10 ppm, and the MS2 mass tolerance was set to 0.6 Da. The false discovery threshold was set strictly to 0.01 using the Percolator Node validated by q-value. The relative abundance of parental peptides was calculated by integration of the area under the curve of the MS1 peaks using the Minora LFQ node.

LFQ was then performed with the Minora feature detector, feature mapper, and precursor ions quantifier nodes. Retention time alignment was performed with maximum RT shift of 5 min and a minimum S/N threshold of 10. Quantified peptides included unique + razor, protein groups were considered for peptide uniqueness, shared Quan results were not used, Quan results with missing values were not rejected, and precursor abundance was based on extracted ion intensity. Imputation was performed using a low abundance resampling method at the peptide level. The Quantitative proteomics data was exported from ProteomeDiscoverer as an excel file and the MitoCoP database (64) was used to remove all non-mitochondrial proteins from the dataset prior to statistical analysis using the R statistical computing environment (R version 4.0.2). Protein abundances were normalized between samples with a random forest (R randomForest version 4.6-14) regression (65), then modeled for expression differences (R limma version 3.44.3) using the linear model fit, with fold changes calculated as a log2 difference. Proteins and peptides identified are listed in Supplementary Table 2.

Volcano plot figures were generated in Python using the matplotlib package. Figures that specify submitochondrial localization are based on annotations from Mitocarta 3.0 (66), with the following modifications: subcellular localizations for TDRKH, HSDL1, and ARMC1 were added manually based on published data (67) (68), TMEM11 is annotated as IM in Mitocarta 3.0; however, a recent preprint provides evidence that TMEM11 is largely in the OM, so this gene was included in analysis of OM proteins in our proteomic data (69).

### Photo-crosslinking of recombinant substrate with isolated mitochondria

An insertion reaction was prepared in a buffer containing 100 mM KoAc, 50 mM HEPES pH 7.4, 2 mM MgoAc_2_, 0.5 mM CaCl_2_. Percoll-gradient enriched mitochondria were added to a final protein concentration of 0.2 mg/mL and CaM-3C-Alfa-Sec61β-OMP25(BpA) was added to a final concentration of 1 µg/mL. Insertion was initiated by the addition of 2 mM EGTA. The insertion reaction was split into -UV and +UV samples. The +UV sample was transferred to a prechilled 6-well plate and left on top of an ice-cooled aluminum block ∼8 cm under a UVP B-100 lamp for 10 minutes.

Both samples were mixed with 10x volumes of IP buffer containing 100 mM KoAc, 50 mM HEPES pH 7.4, 2 mM MgoAc_2_, and 1% Triton X-100. After incubating on ice for 10 minutes, samples were centrifuged at 18,000 rpm. Supernatant was mixed with 10 µL packed streptavidin agarose resin (Thermo Scientific Cat. #20357) which had been functionalized with biotinylated Alfa-Nb, blocked with free biotin, and equilibrated in IP buffer. After 1 hour of binding head-over-tail at 4°C, the unbound fraction was removed, and the resin was then washed 2x with 1 mL IP wash buffer, 2x with 1 mL IP wash buffer + 0.5 M NaCl, and 2x with 1 mL IP wash buffer. Resin was then incubated with 10 µL IP wash buffer + 300 nM SUMOstar protease (LifeSensors Cat # SP4110) for 30 minutes on ice. 1 mL IP wash buffer was added to protease treated resin and incubated for 5 minutes on ice. Eluted protein was then removed from resin and TCA precipitated by adding 1:10 volume 100% TCA and incubating on ice for 10 minutes, followed by centrifuging at max speed in a chilled benchtop centrifuge for 10 minutes. The pellet was then washed 2x with ice-cold acetone and prepared for mass spectrometry as described below.

Samples used for western blotting were prepared with the following differences: isolated mitochondria were not purified on a percoll gradient; pierce magnetic streptavidin resin was used (Thermo Scientific Cat. #88817), an IP buffer containing 200 mM NaCl, 50 mM HEPES pH 7.4, 1% Triton X-100 was used for solubilization; 4x 1 mL wash steps; and an elution buffer containing 200 mM NaCl, 50 mM HEPES pH 7.4, and 0.05% Triton X-100 was used for 2x wash steps and then elution was performed in the same buffer + 300 nM SUMOstar in a 10 µL volume.

### Proteoliposome reconstitutions and insertions

Reconstitutions of protein into liposomes were similar to previously described methods (70) (26). The following phospholipids were obtained from Avanti Polar Lipids: phosphatidyl-choline (PC) and phosphatidyl-ethanolamine (PE) from egg, and synthetic 1,2-dioleoyl-sn-glycero-3-phosphoethanolamine-N-lissamine rhodamine B (Rh-PE). The liposome mixture contained PC:PE:Rh-PE at a mass ratio of 8:1.9:0.1. Rh-PE was used to monitor recovery throughout the reconstitutions and for quantification. Lipids were mixed at the indicated ratio as chloroform stocks, adjusted to 10 mM DTT and dried by centrifugation under vacuum overnight. The resulting lipid film was rehydrated to a final concentration of 20 mg/mL in lipid buffer (15% glycerol, 50 mM HEPES pH 7.4) and mixed for 8 hours at 25°C until a homogenous mixture was achieved. Lipids were then diluted with more lipid buffer and supplemented with DBC to produce a lipid/DBC mixture containing 2% DBC and 10 mg/mL lipids. BioBeads-SM2 (Bio-Rad) were prepared by activation with methanol, washing thoroughly with distilled water, and then were resuspended in water into a final slurry where they occupied 50% of the final volume. For reconstitutions, excess liquid was removed from BioBeads by aspiration just before use. Reconstitutions used purified EMC and MTCH2 in 0.25% DBC were obtained as described above. In initial experiments, we determined the relative concentration of purified MTCH2 compared to the amount in isolated mitochondria from K562 cells. Different dilutions of purified MTCH2 were mixed with constant amounts of lipids and adjusted to a final buffer concentration of 100 mM NaCl, 25 mM HEPES pH 7.4, 2 mM MgCl_2_, 0.8% DBC. Liposomes were made using the same buffer and detergent conditions. A standard 100 µL reaction contained 10-40 µL purified MTCH2, 20 µL of the 10 mg/mL lipid/DBC mixture, and the remaining volume made up with buffer, salts, and detergent. This protein/lipid/detergent mixture was added to 120 µL BioBeads in 1.5 mL Eppendorf tubes. The slurry was mixed in a thermomixer for 18 hours at 4°C. The fluid phase was removed and to another 1.5 mL Eppendorf tube containing 120 µL Biobeads, and mixed in a thermomixer for 2 hours at 23°C. The fluid phase was then separated and diluted with ten volumes of ice-cold water. The proteoliposomes were sedimented in a TLA120.2 rotor at 70,000 rpm for 30 minutes, and resuspended in 16.7 µL liposome resuspension buffer (100 mM KoAc, 50 mM HEPES pH 7.4, 2 mM MgoAc2, 250 mM sucrose, 1 mM DTT). Substrates for insertions were prepared by translating in RRL for 15 min, then the reactions were treated with 1 mM puromycin to prevent further synthesis. Insertion reactions consisted of 8 µL translation in RRL, 1 mM EGTA and 2 µL buffer, liposomes, proteoliposomes, or isolated mitochondria. The reactions were incubated at 32°C for 30 min, before being treated with 0.5 mg/mL PK for 1 hour on ice. The reactions were quenched and the protected fragments enriched with NiNTA resin as described above.

### Flow cytometry

For all reporter experiments, respective K562 cell lines were spinfected with lenti-virus for indicated constructs and analyzed by flow cytometry after 48-72 hours. All reporter experiments were performed at least twice. For the apoptosis experiment, wildtype K562 cells expressing either MTCH2-P2A-BFP or BFP alone were treated with 5 μM imatinib mesylate (461080010, Fisher Scientific, USA) for 72 hours. Cells were harvested, washed once with ice cold PBS, then resuspended in 100 μL staining buffer (10 mM HEPES, 140 mM NaCl, 2.5 mM CaCl2 pH 7.4, 5% Annexin-FITC (Invitrogen, A13199), 50 μg/mL propidium iodide (P1304MP, Invitrogen, USA) before analysis by flow cytometry. For treatment with the GPAT inhibitor FSG67, K562 cells were treated for 16 hours with 75 μM FSG67 (Cedarlane Laboratories Cat. #10-4577) as previously described (19) prior to analysis by flow cytometry. All samples were run on either an NXT Flow Cytometer (ThermoFisher) or a MACSQuant VYB (Mitenyi Biotec). Flow cytometry data was analyzed either in FlowJo v10.8 Software (BD Life Sciences) or Python using the FlowCytometryTools package.

### Microscopy

To visualize localization to the mitochondria, K562 cells were stained with 1 μM Mitotracker Deep Red FM (M22426, ThermoFisher) for 30 minutes. Cells were collected, spun down, washed in fresh media and plated onto a 96-well glass bottomed plate (160376, Thermo Fisher). K562 cells were then briefly spun down by centrifugation at 100*g* for 5 minutes and imaged on a Zeiss LSM710 NLO Laser Scanning confocal microscope.

### Sequence alignments

An alignment of individual SLC25 repeats from various human transporter was generated using the hmmalign tool from the HMMER3.3 package (71) and a precomputed hmm profile of the SLC25 family from pfam (pf00153) (72).

---

**Fig S1.**
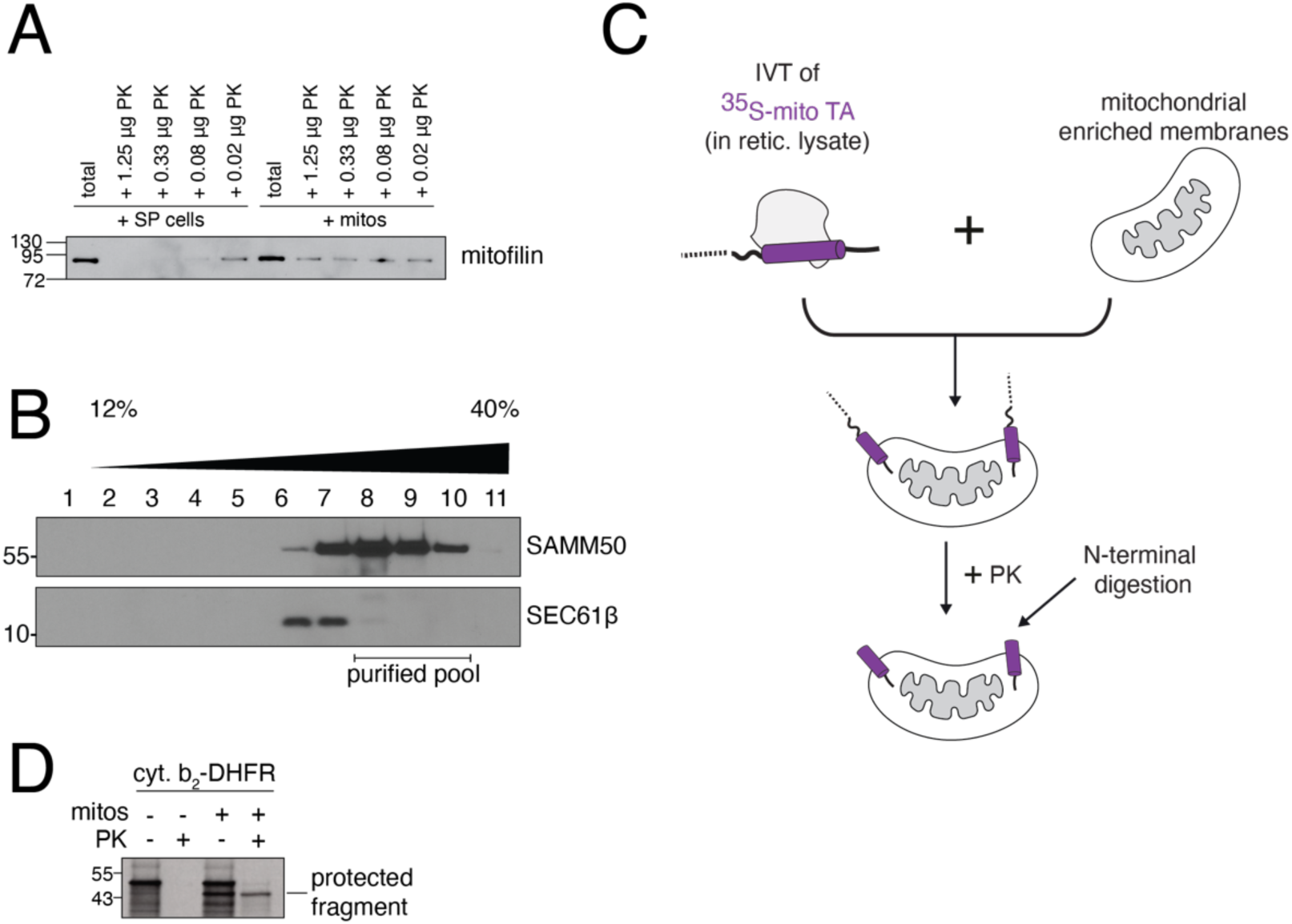
In vitro protease protection assays into human mitochondria. **(A)** The integrity of the outer membrane was tested using both semi-permeabilized K562 cells and mitochondrially enriched membrane fractions by treating each with decreasing amounts of proteinase K (PK). The resulting reactions were analyzed by western blot for the inner mitochondrial membrane protein mitofilin, which should be inaccessible to PK when the outer membrane is intact. **(B)** Further purification of mitochondria from K562 cells using a percoll gradient and density centrifugation. Blots for a mitochondrial marker (SAM50) and an ER marker (SEC61β) demonstrate relative proportion of mitochondria and ER in the ‘mitochondrial enriched membranes’ that are used for most in vitro experiments. Proteomics and crosslinking-mass spectrometry experiments were performed with percoll-gradient enriched mitochondria and used fractions 8-10 (Fig. 2A, Fig. 3C). **(C)** Schematic of the basic in vitro insertion assay, using protease protection as a readout for insertion. In all insertion assays, substrates were translated using an in vitro translation reaction (IVT) in rabbit reticulocyte lysate and ^35^S-methionine labelled, permitting detection by autoradiography. Translation was terminated by addition of puromycin, followed by incubation with membranes enriched in mitochondria derived from human K562 cells. After incubating with membranes, reactions were treated with PK. For TA proteins, the resulting protease protected band was immunoprecipitated via a tag on its C-terminus, ensuring insertion into the outer membrane in the correct orientation, and analyzed by SDS-PAGE and autoradiography. **(D)** The IMS-localized TOM substrate composed of the cytochrome b targeting sequence fused to the inert protein DHFR (cyt. b_2_-DHFR) was translated and incubated with purified mitochondria to confirm their activity. Insertion was detected by protease protection as described.

**Fig S2.**
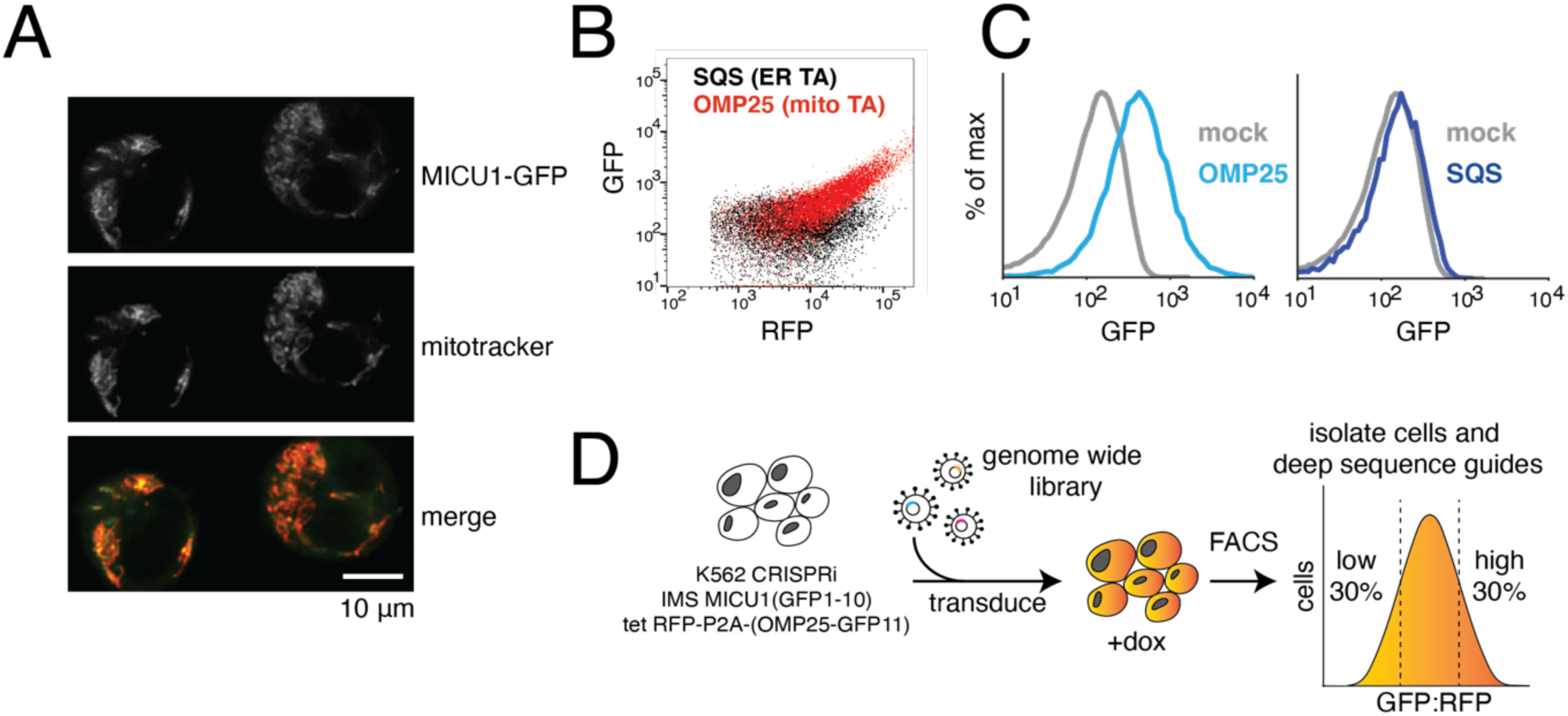
A CRISPRi screening platform to identify factors involved in mitochondrial TA protein biogenesis in human cells. **(A)** Microscopy showing that the IMS-targeting sequence from MICU1 conjugated to full length GFP results in its localization to the mitochondria in mammalian K562 CRISPRi cells. **(B)** The endogenous sequences of two TA proteins: SQS, which is localized to the ER, and OMP25, which under these conditions is dual-localized to both the outer membrane and ER, were appended to a C-terminal GFP11 in a backbone containing a translational control (RFP) separated by a viral 2A sequence (see Fig. 1B). These constructs were independently introduced into cells expressing IMS localized GFP1-10 and analyzed by flow cytometry. **(C)** Histograms of (B) comparing GFP fluorescence for OMP25 and SQS compared to a mock transduced control. The marked increase in GFP fluorescence suggest that OMP25, but not SQS can successfully conjugate with GFP1-10 localized to the IMS. **(D)** Workflow of the FACS-based CRISPRi screen. A K562 CRISPRi reporter cell line was constructed that constitutively expressed GFP1-10 in the IMS and the OMP25-GFP11 reporter under an inducible promoter. For the screen, these cells were transduced with a genome-scale CRISPRi sgRNA library and then the OMP25-GFP11 reporter was induced with doxycycline for 24 hours prior to cell sorting. Cells were sorted based on ratiometric changes in GFP relative to RFP, and sgRNAs expressed in the isolated cells were identified using deep sequencing.

**Fig S3.**
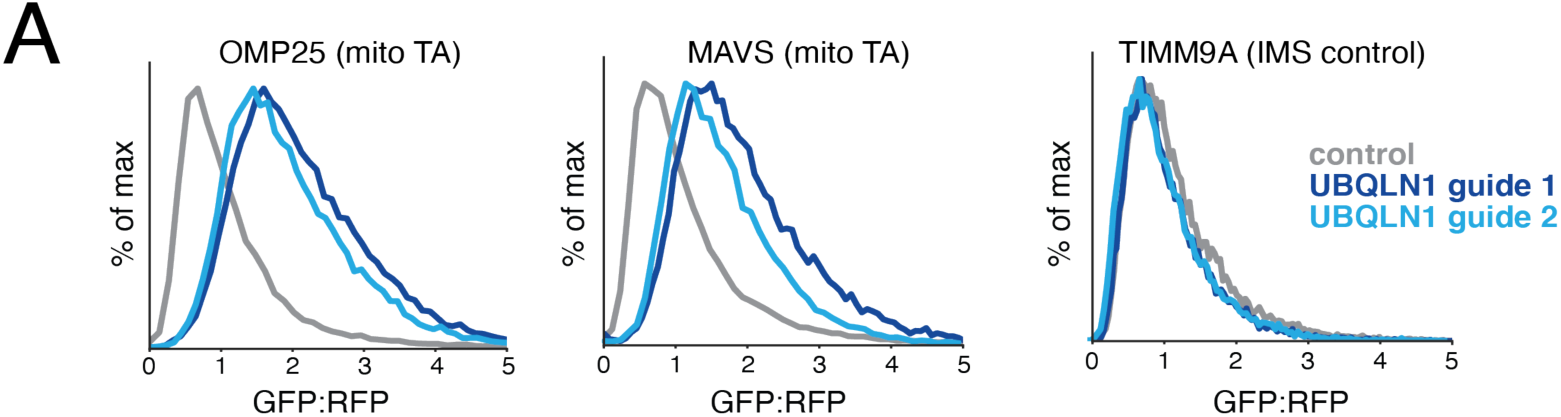
UBQLN1 is a quality control factor for mitochondrial TAs. K562 CRISPRi cells expressing IMS GFP1-10 were depleted of UBQLN1 using two different sgRNAs. Reporters were introduced for either two mitochondrial TAs (OMP25 and MAVS) or an IMS localized control (TIM9A) and cells were analyzed by flow cytometry. Lack of UBQLN results in a ratiometric increase in GFP:RFP fluorescence for mitochondrial TAs, consistent with its previously reported role in targeting mislocalized mitochondrial TAs for degradation by the ubiquitin proteasome pathway (16).

**Fig S4.**
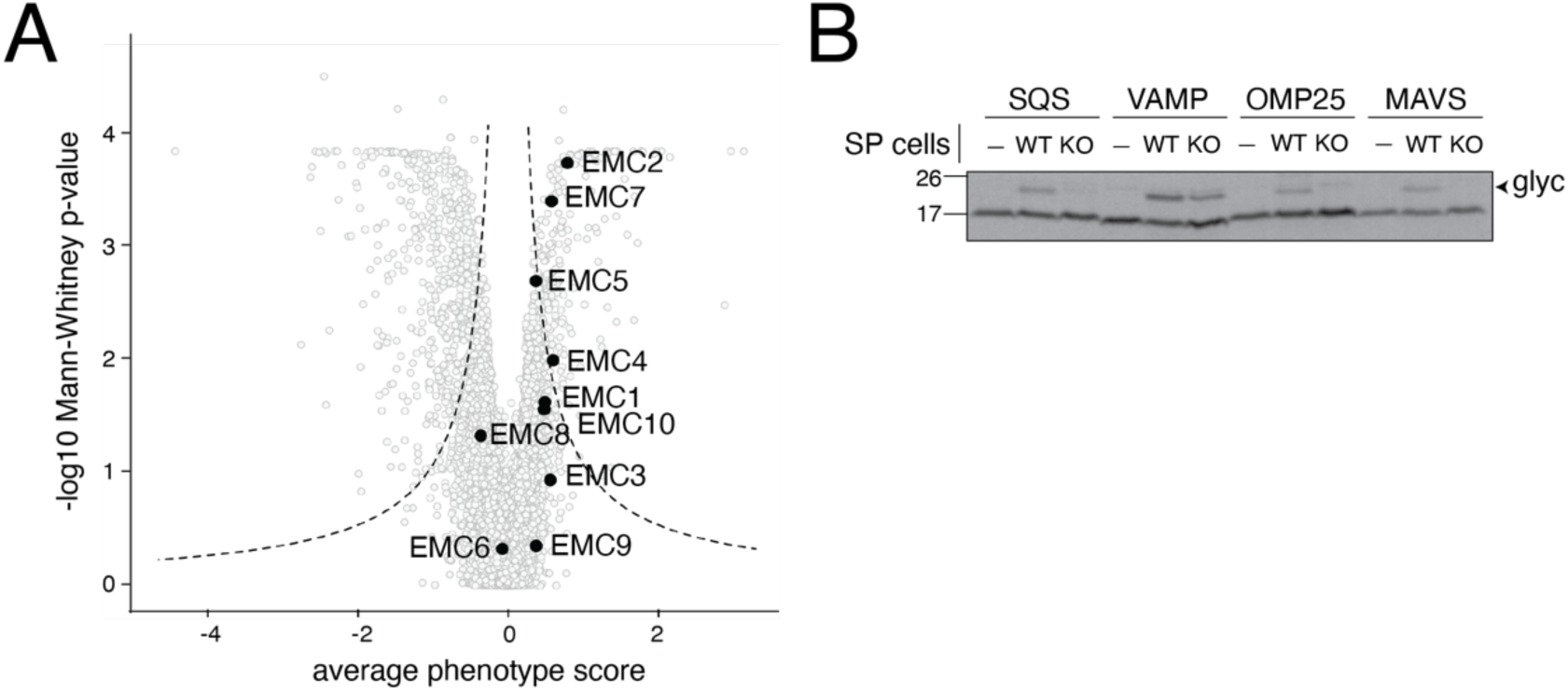
The EMC is required for insertion of mislocalized mitochondrial TAs to the ER. **(A)** Volcano plot of the genome-wide CRISPRi screen with EMC subunits shown in black. **(B)** A panel of ER (SQS and VAMP) and mitochondrial (OMP25 and MAVS) TA proteins were conjugated to a C-terminal opsin epitope. The opsin epitope contains a consensus glycosylation sequence that is modified upon insertion into the ER lumen. These constructs were then translated in rabbit reticulocyte lysate in the presence of ^35^S-methionine. The reactions were puromycin treated and incubated with either wild-type (WT) or EMC knockout (KO) semi-permeabilized (SP) cells. Insertion into the ER, as monitored by appearance of a glycosylated band (‘glyc’), was dependent on the EMC for its canonical substrate SQS, and both mitochondrial TAs. In contrast, VAMP’s insertion was unaffected by EMC knockout, consistent with its previously reported dependence on the GET pathway for insertion (26).

**Fig S5.**
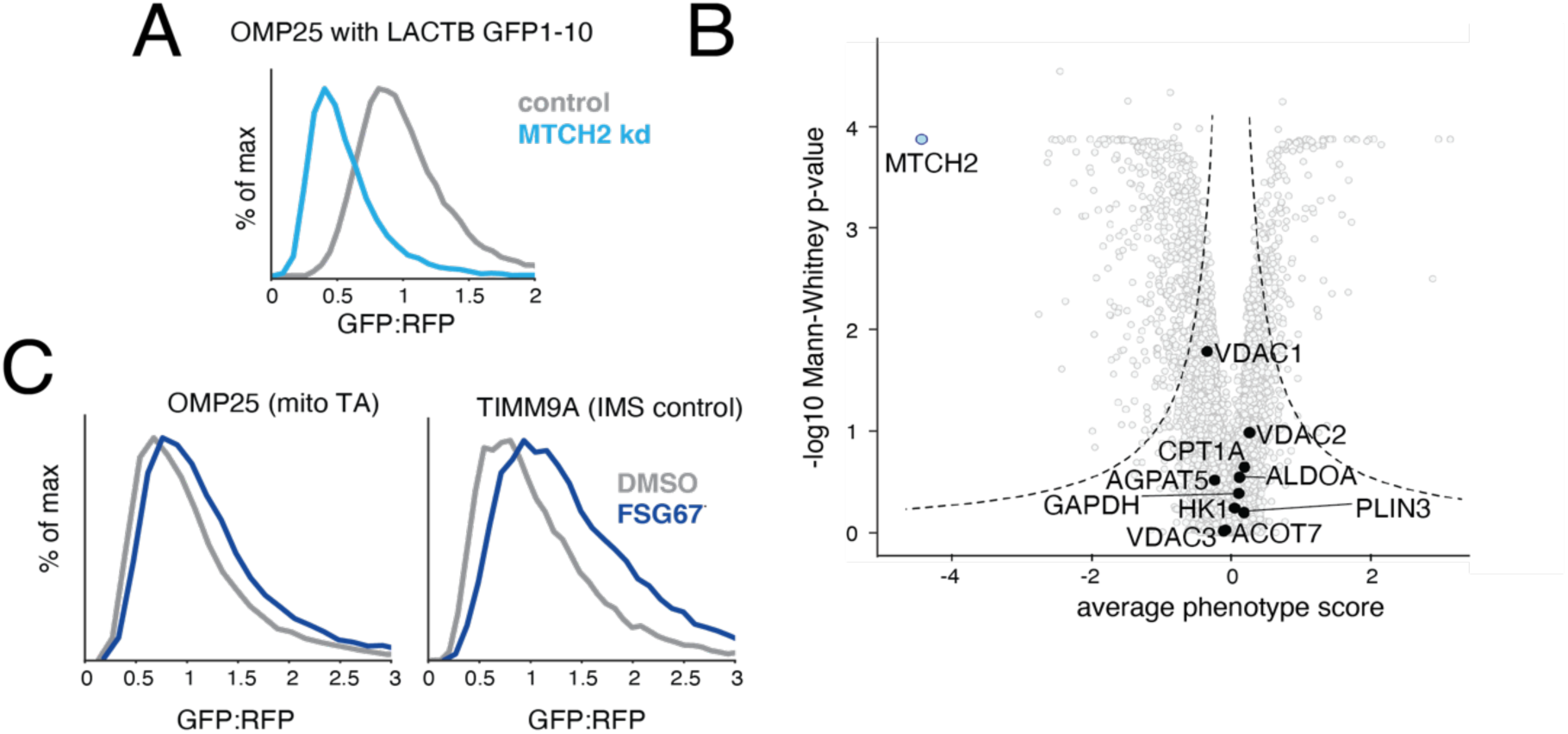
Assessing the effects of lipid biogenesis defects on mitochondrial TAs. **(A)** Flow cytometry analysis as in Fig. 1D but with an alternative IMS targeting sequence derived from LACTB appended to the GFP1-10 (44). (**B**) Volcano plot of the genome-wide CRISPRi screen indicating MTCH2 (in light blue) and factors previously implicated in the regulation of outer membrane fatty acid synthesis or transport (in black). **(C)** K562 IMS GFP1-10 expressing cells were treated with the pan GPAT inhibitor FSG67 for 16 hours (75 μM) or a vehicle. MTCH2-dependent mitochondrial fusion has been shown to require Glycerol 3-phosphate acyltransferases (GPATs) catalysed LPA synthesis. A reporter expressing either a mitochondrial TA (OMP25) or an IMS localized control (TIM9A) were expressed in GPATi cells and analyzed by flow cytometry.

**Fig. S6.**
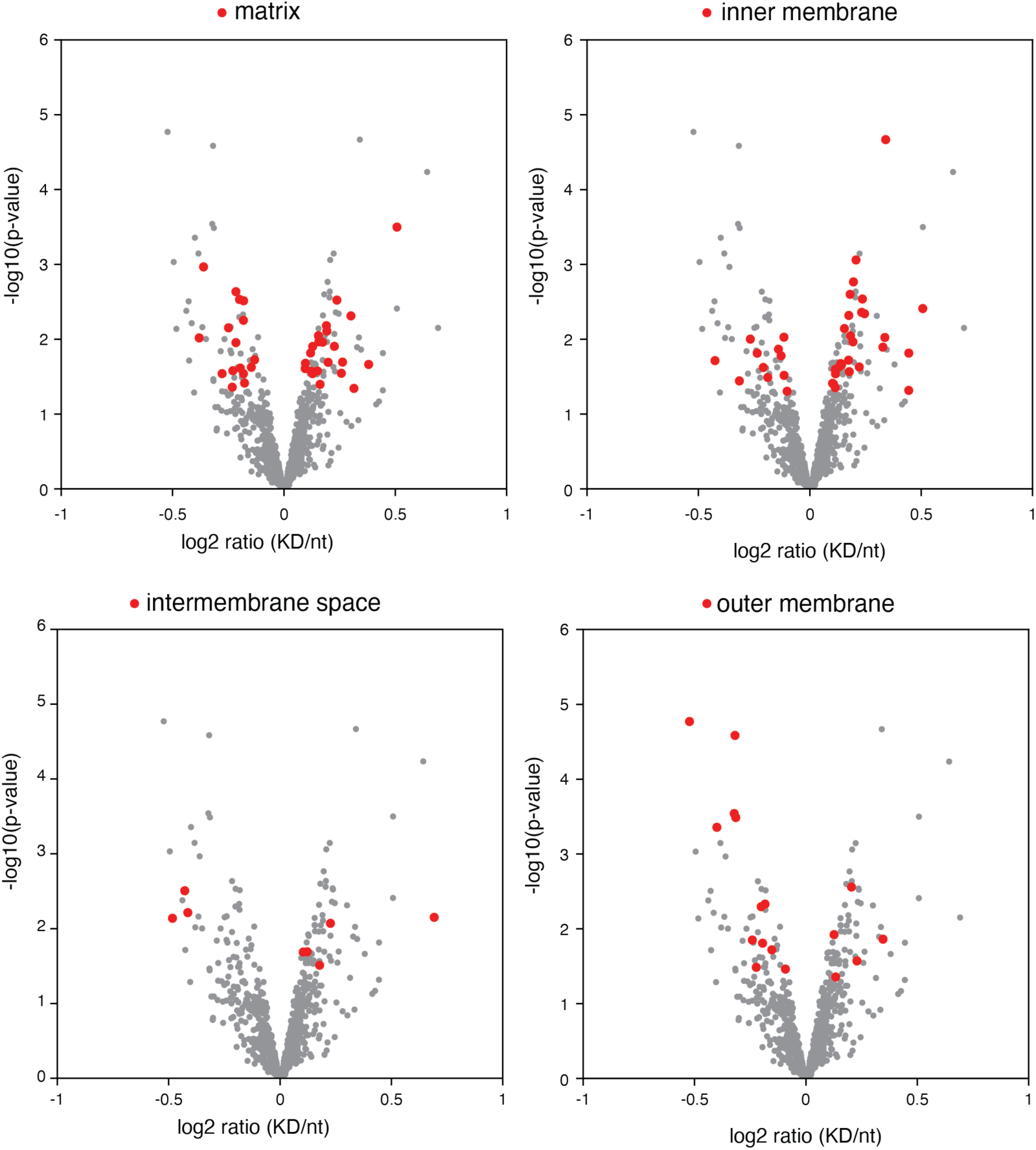
MTCH2 depletion affects endogenous outer mitochondrial membrane proteins. As in Fig. 2A, but with all proteins from the indicated mitochondrial compartment which have a p-value > 0.05 highlighted in red.

**Fig. S7.**
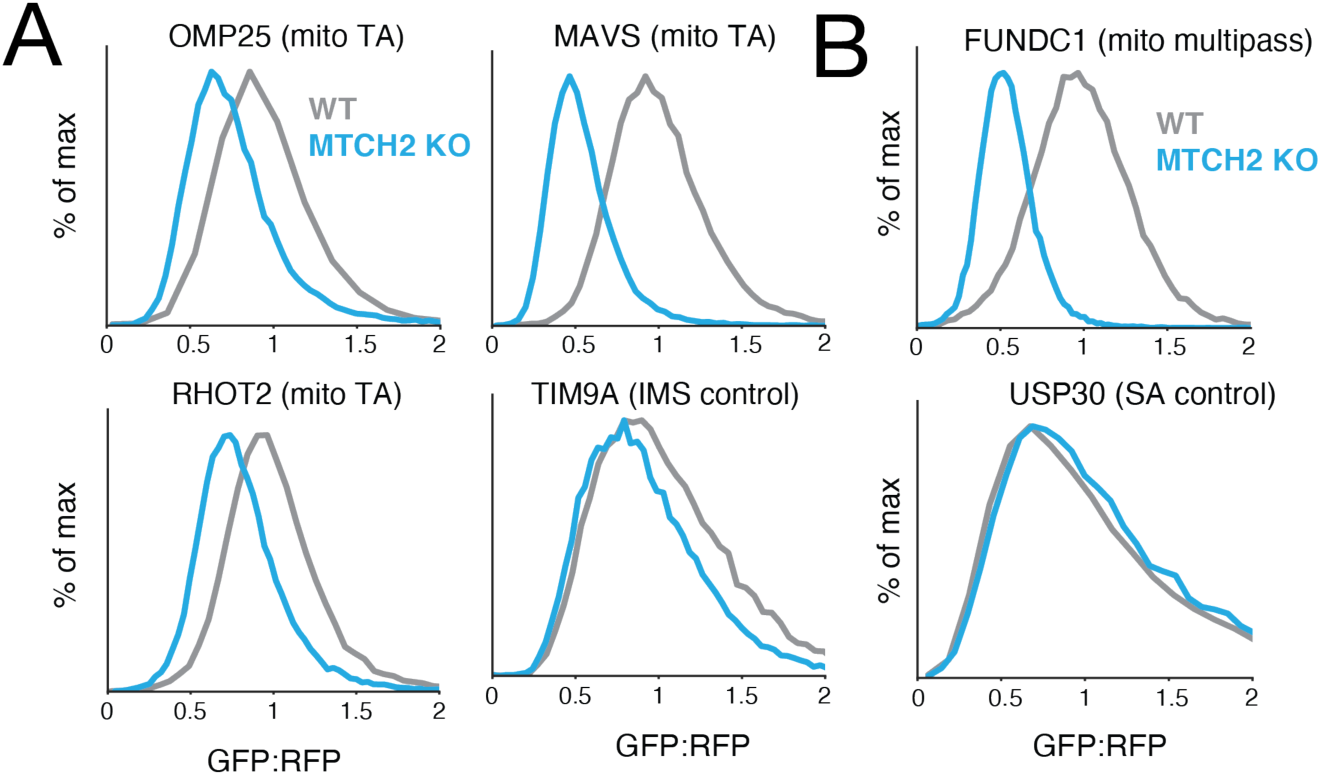
Analysis of TA proteins for MTCH2 dependent biogenesis in knockout cells. As in Fig. 2C but in wild type compared to MTCH2 knockout cells. Histograms summarizing flow cytometry analysis of the integration of the indicated mitochondrial proteins.

**Fig. S8.**
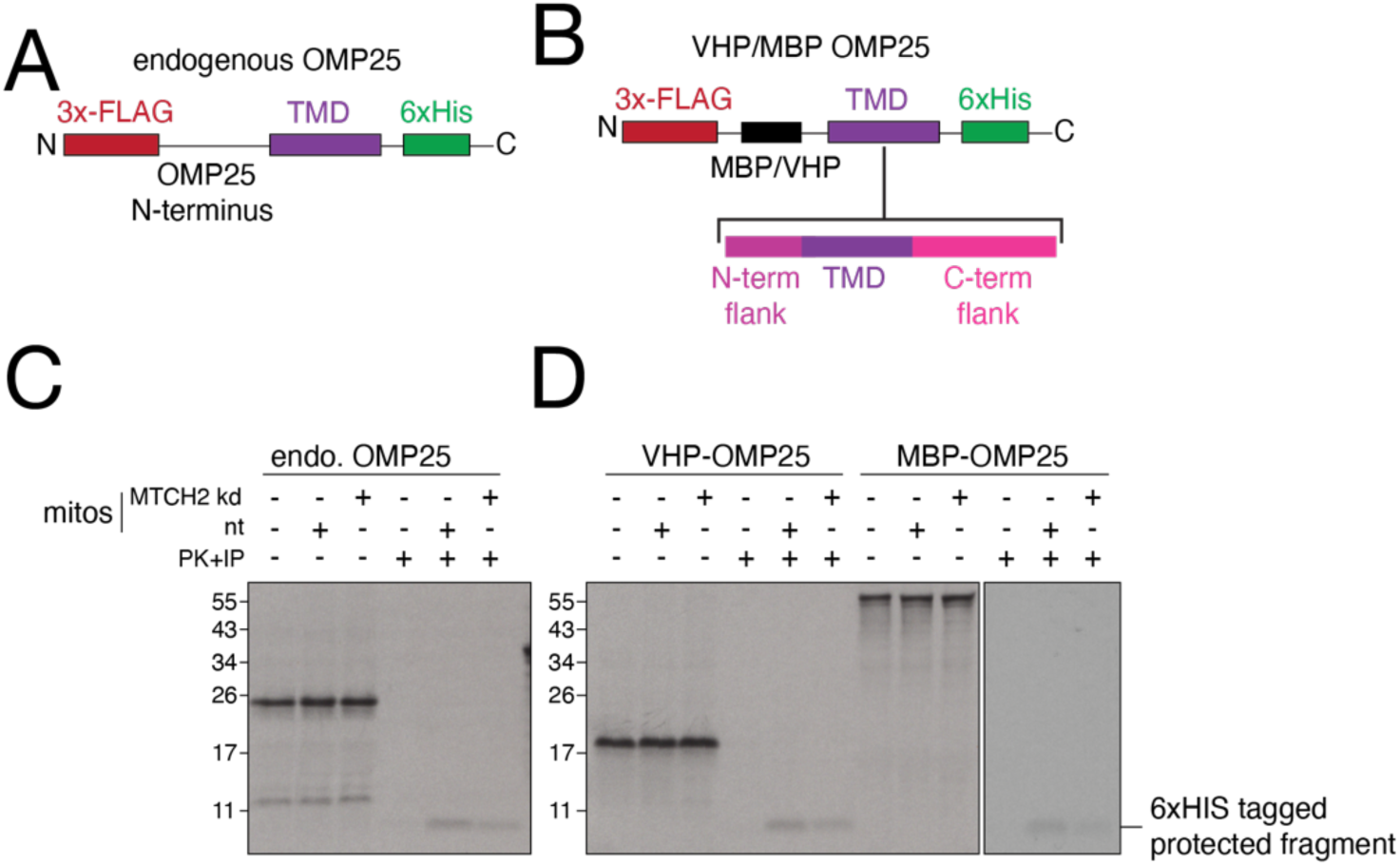
Establishing an in vitro system to test differential dependence on MTCH2 on insertion of a panel of mitochondrial TAs. **(A, B)** Schematic of OMP25 constructs used to test whether the dependence on MTCH2 for insertion that we observed with the full-length endogenous OMP25 (A) could be recapitulated with an artificial N-terminus (B). **(C)**Using the protease protection assay described in fig. S1, we compared insertion of the endogenous OMP25 containing a C-terminal 6xHIS tag into mitochondria isolated from K562 cells expressing either a non-targeting (nt) or MTCH2 sgRNA (kd). We observed that loss of MTCH2 specifically decreased the levels of a protease protected fragment consistent in size with the TMD and C-terminus of OMP25 (compare lanes 5 vs 6). In the absence of mitochondria, no protected fragment is observed (lane 4). Because this protease protected fragment could be immunoprecipitated using a C-terminal 6xHIS tag, we verified that OMP25 was inserted in the correct orientation, with its C-terminus in the IMS and its N-terminus facing the cytosol. **(D)** As in (A) but using a fusion of the OMP25 TMD and its flanking residues to both the unrelated globular proteins MBP and VHP. We concluded that the OMP25 TMD alone is sufficient to confer MTCH2 dependent insertion on both of the tested fusion proteins (compare lanes 5 vs 6 and 11 vs 12). Because the VHP-fusions were translated more efficiently, we generated a panel of mitochondrial TAs using the depicted VHP N-terminus as shown in (B).

**Fig. S9.**
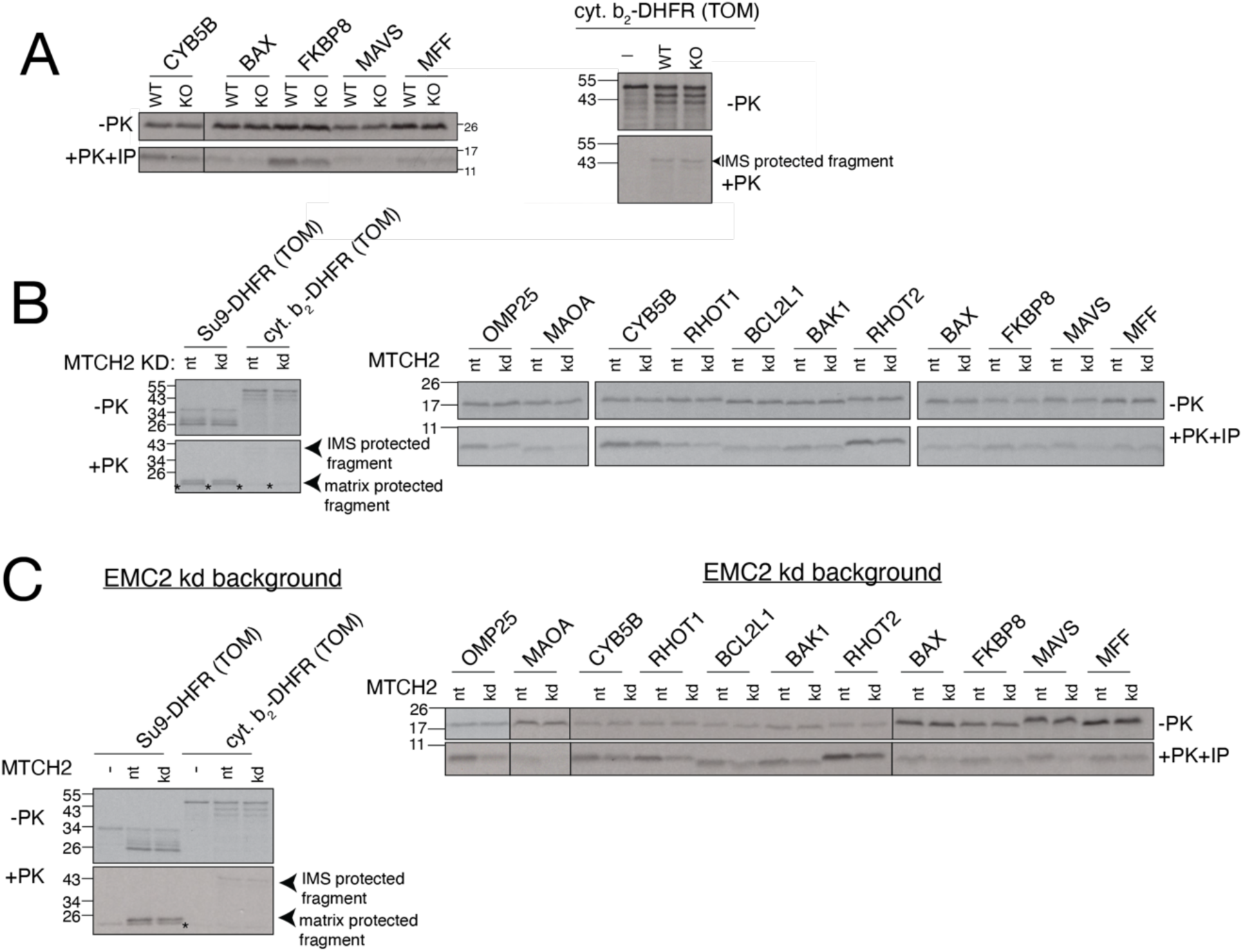
Insertion of mitochondrial TAs in vitro is affected by MTCH2 knockdown. **(A)** Additional substrates were tested in parallel with the data displayed in Fig. 3A, where the indicated TMDs and flanking residues were fused to VHP and examined for insertion into wild type (WT) and MTCH2 knockout (KO) mitochondria. As a control, a canonical IMS localized TOM substrate derived from a fusion of the cytochrome b targeting sequence to DHFR (cyt. b_2_-DHFR) was tested in parallel. **(B)** As in Fig 3A except using mitochondria isolated from K562 CRISPRi cells expressing either a non-targeting (nt) or MTCH2 targeting (kd) sgRNA. A panel of mitochondrial TAs was tested in parallel with a matrix (Su9-DHFR) and IMS (cyt. b_2_-DHFR) targeted control that rely on the TOM pathway, which were unaffected. *Denotes the folded, protease resistant, DHFR domain that migrates immediately below the mature, matrix targeted control, and is visible in the absence of mitochondria. **(C)** Note that because some residual ER is present in the enriched mitochondrial membranes used in these insertion reactions (see fig. S1), for those substrates that are dual localized, we cannot formally differentiate between insertion into mitochondria vs ER in this assay alone. To address this, we have tested the complete TA panel in an EMC knockdown background, which we found eliminated mistargeting of mitochondrial TAs to the ER (fig. S4). Therefore, we perform insertion assays as in (A) except using (see fig. S1B) mitochondria isolated from either EMC2 knock down or EMC2 and MTCH2 knock down K562 CRISPRi cells. The insertion defect in the absence of MTCH2 was enhanced for some substrates (see CYB5B) when entry to the ER was blocked, suggesting these substrates may be partially dual localized under these conditions.

**Fig. S10.**
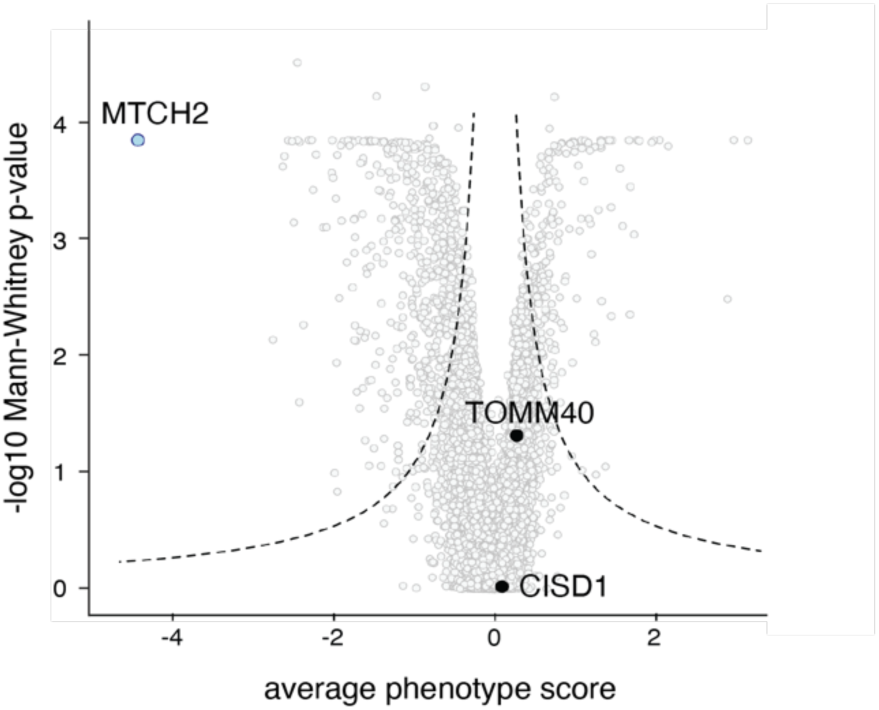
TOM40 and CISD1 are not significant hits in a mitochondrial TA biogenesis CRISPRi screen. Volcano plot of the genome-wide CRISPRi screen as in Fig. 1C indicating MTCH2 (in light blue) and the two other factors (TOM40 and CISD1, in black) that were enriched upon crosslinking OMP25 with purified mitochondria (Fig. 3C; see fig. S1C for mitochondrial purification).

**Fig. S11.**
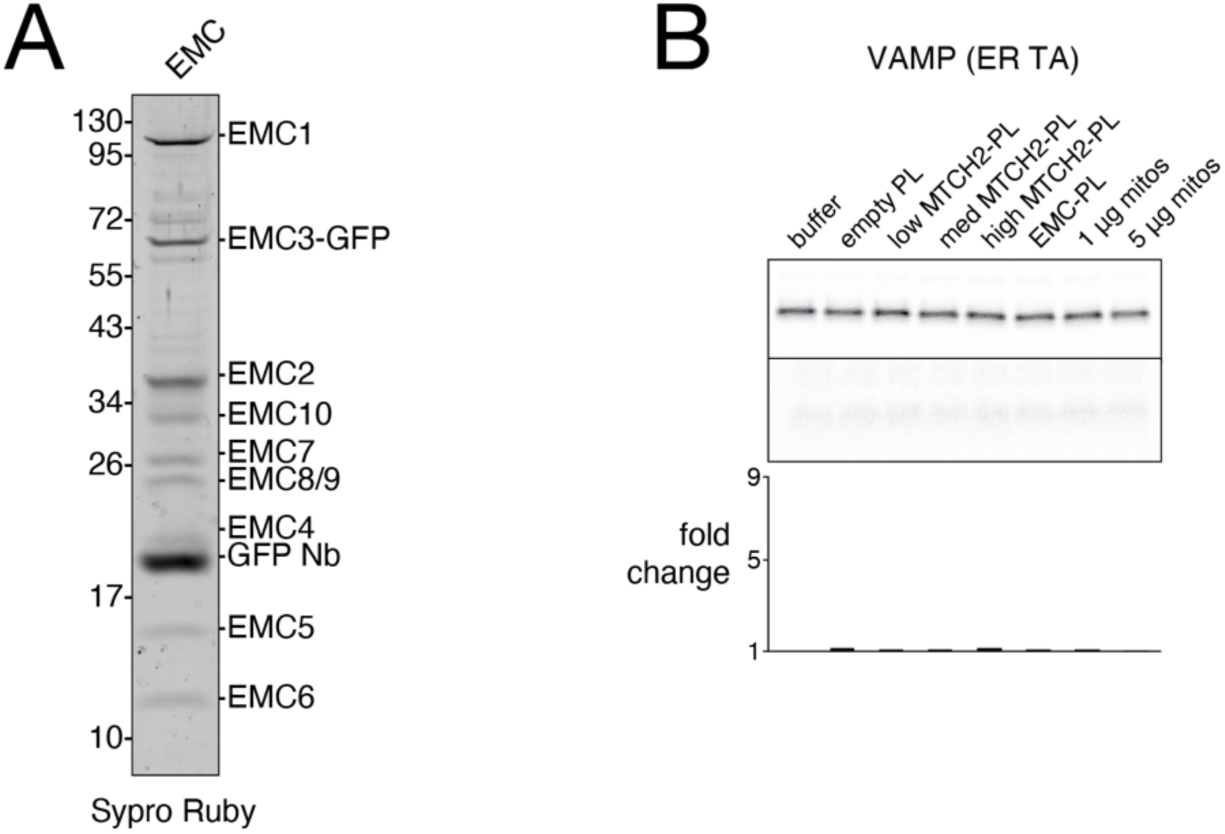
Purification of the human EMC and ER TA insertion control. **(A)** The human EMC was expressed and purified as previously described using a GFP fused to the C-terminus of EMC3 in the detergent DBC (25) and visualized using Sypro Ruby staining. This was further used for generating EMC-proteoliposomes as indicated in Fig. 3E-F. (B) As in Fig. 3F but with an ER TA protein (VAMP) that is dependent on the GET-pathway for insertion.

**Fig S12.**
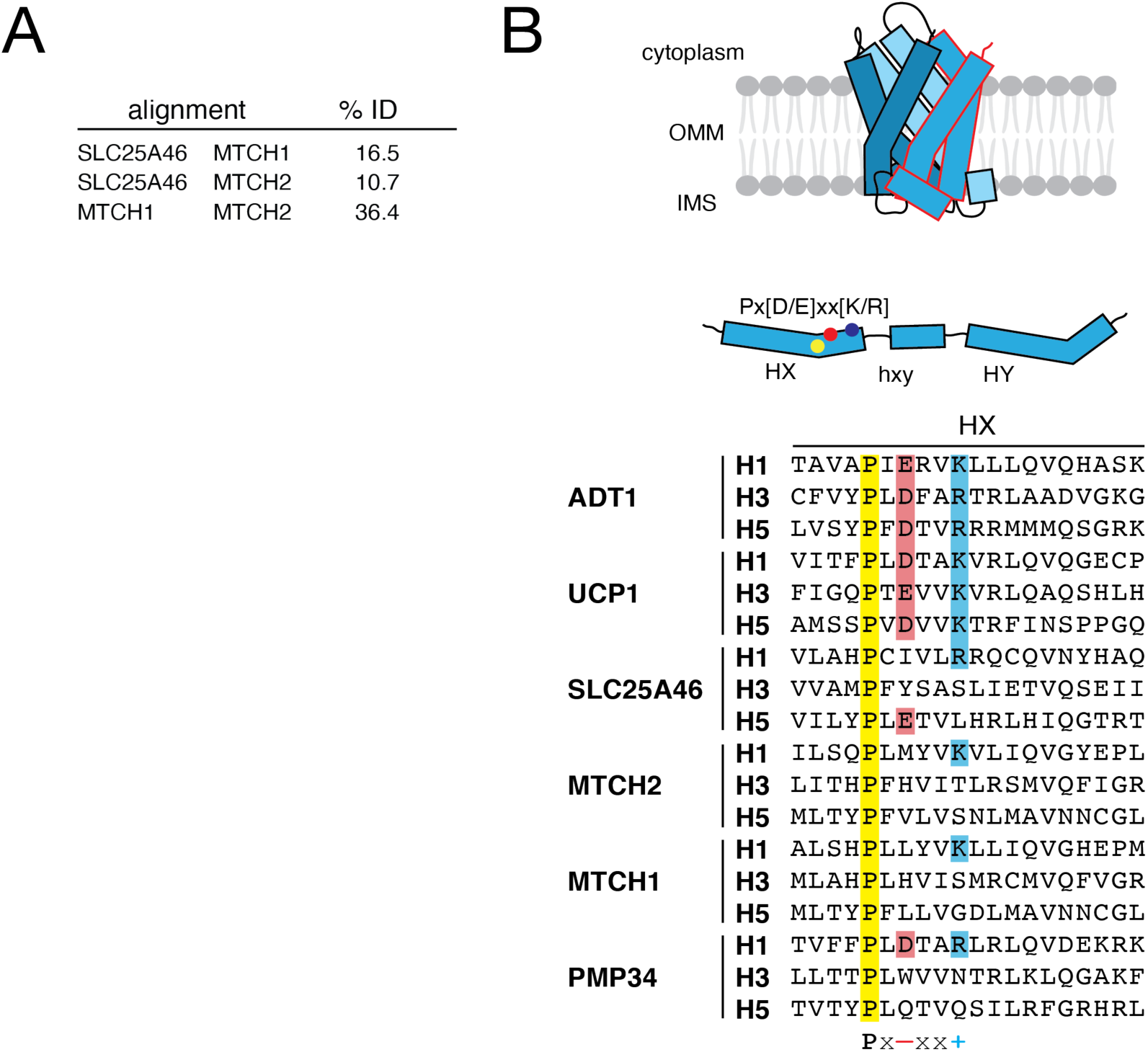
Shared features of outer membrane SLC25 transporters. **(A)** Sequence identity derived from pairwise alignment between SLC25A46, MTCH1, and MTCH2. **(B)** On top, a cartoon representation of SLC25 TMD arrangement showing the 3 SLC25 repeats in unique shades of blue, with a single repeat outlined in red. In the middle, a schematic showing the location of characteristic motifs within a single SLC25 repeat, which normally encodes 2 TM helices. On bottom, sequence alignment of all individual SLC25 repeats from 2 inner membrane SLC25 transporters (ADT1, UCP1) and 4 outer membrane SLC25 transporters, three mitochondrial (SLC25A46, MTCH1, MTCH2) and one peroxisomal (PMP34), with residues from the Px[D/E]xx[K/R] motif highlighted.

**Fig. S13.**
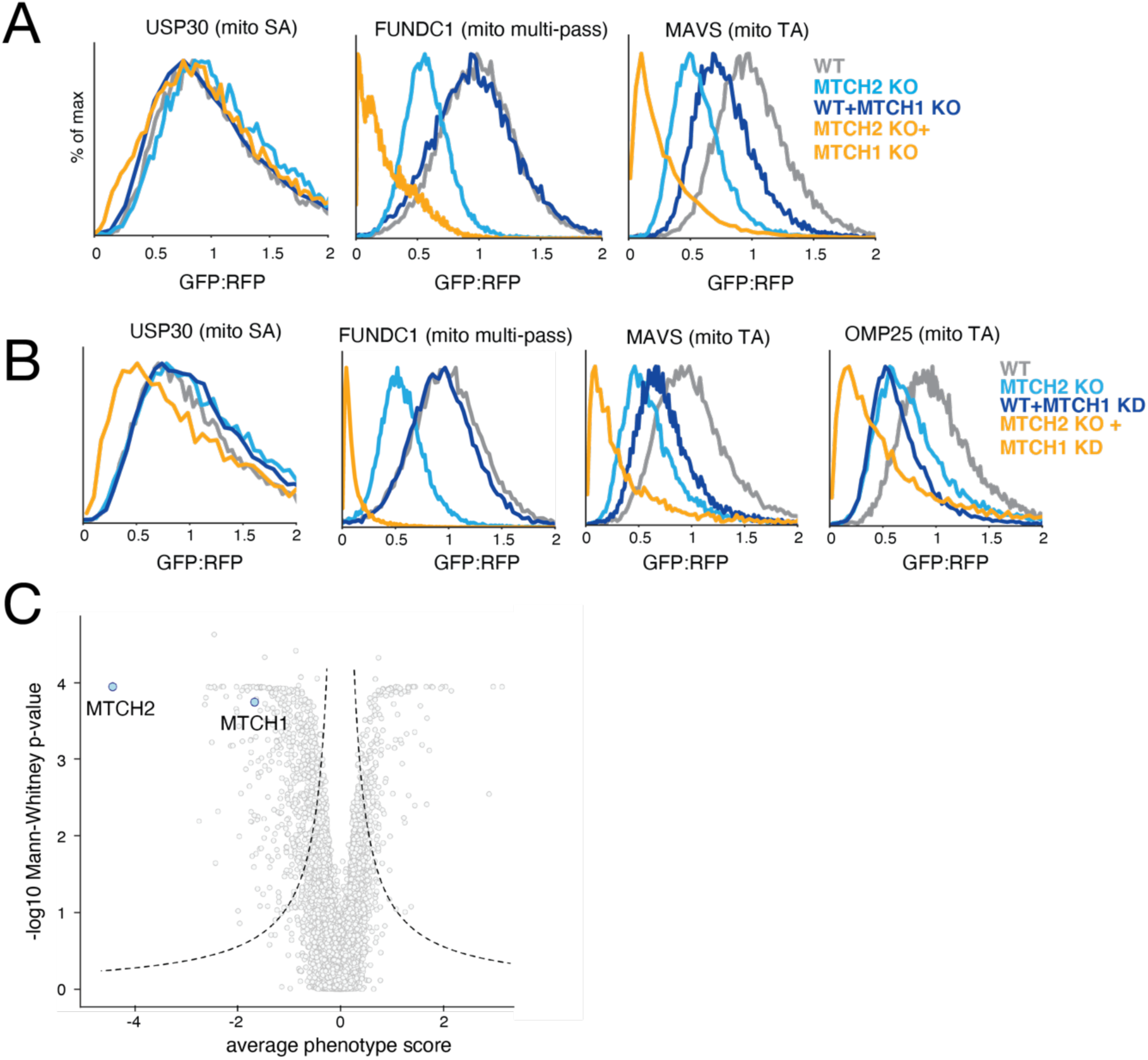
MTCH1 acts in a parallel pathway to mediate insertion of mitochondrial TAs. **(A)** As in Fig. 4B for a set of outer membrane reporters including a signal-anchored protein (USP30), a mitochondrial TA (MAVS) and a multipass protein (FUNDC1). **(B)** As in (A) but in either wild-type or MTCH2 KO K562 CRISPRi cells expressing guides targeting MTCH1 for knock-down. **(C)** Volcano plot of the genome-wide CRISPRi screen with MTCH1 and MTCH2 highlighted.

**Fig. S14.**
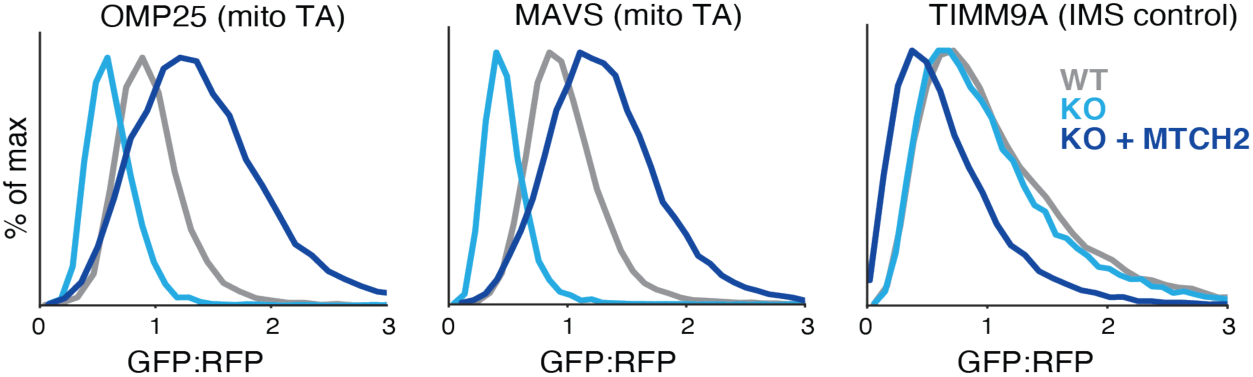
MTCH2 overexpression drives insertion of mitochondrial TAs into the outer membrane. Flow cytometry analysis of two mitochondrial TAs (OMP25 and MAVS) and an IMS control (TIM9A) in the indicated cell types.

**Fig. S15.**
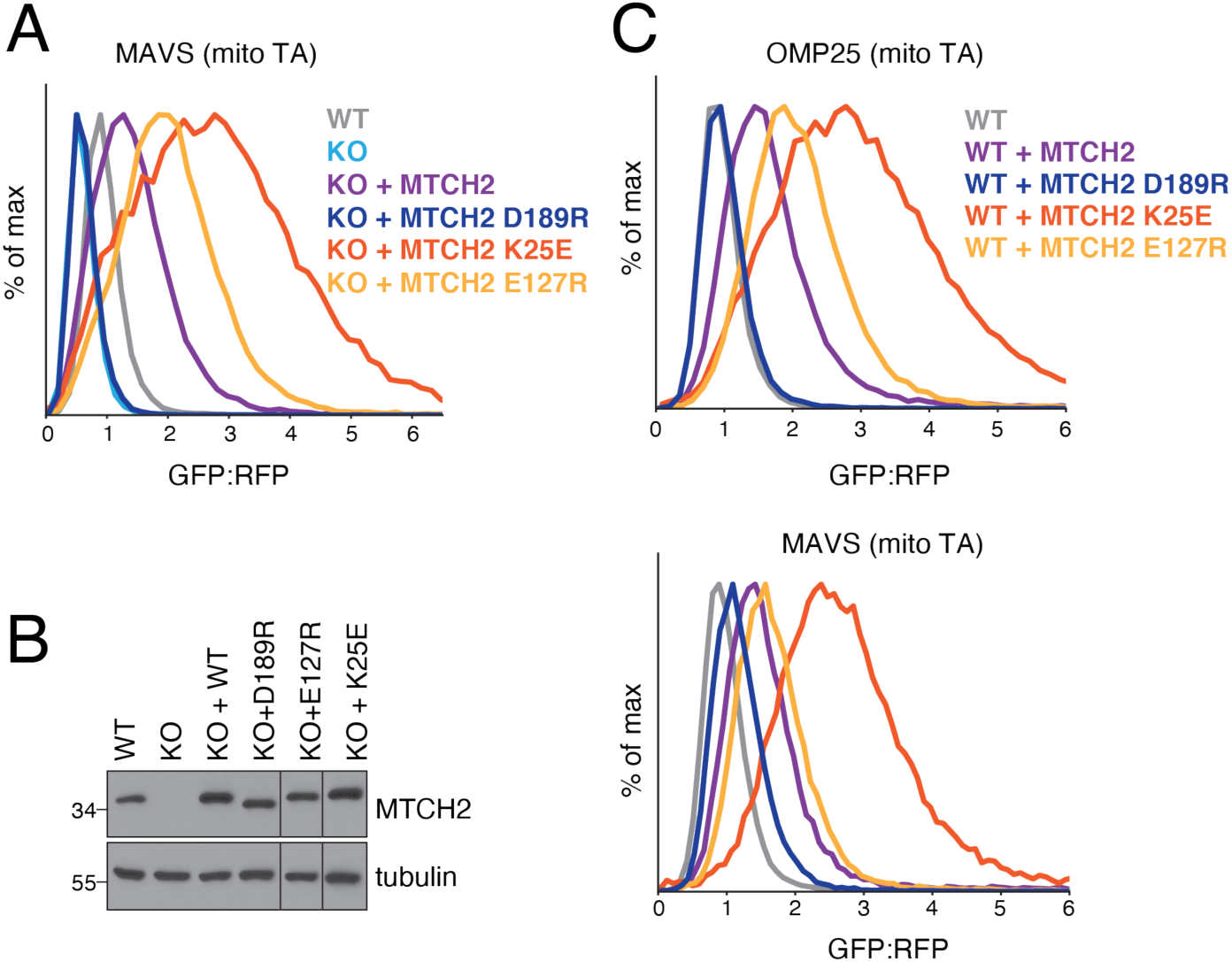
Point mutants to MTCH2 TMDs affect integration of mitochondrial TAs. **(A)** As in Fig. 4C for another mitochondrial TA (MAVS). **(B)** Blots showing expression of wild-type MTCH2 and the indicated point mutants in a K562 MTCH2 KO background relative to wild-type cells. **(C)** As in (A) except point mutants were expressed on top of wild-type (WT) cells which are expressing functional copies of MTCH2.

**Fig. S16.**
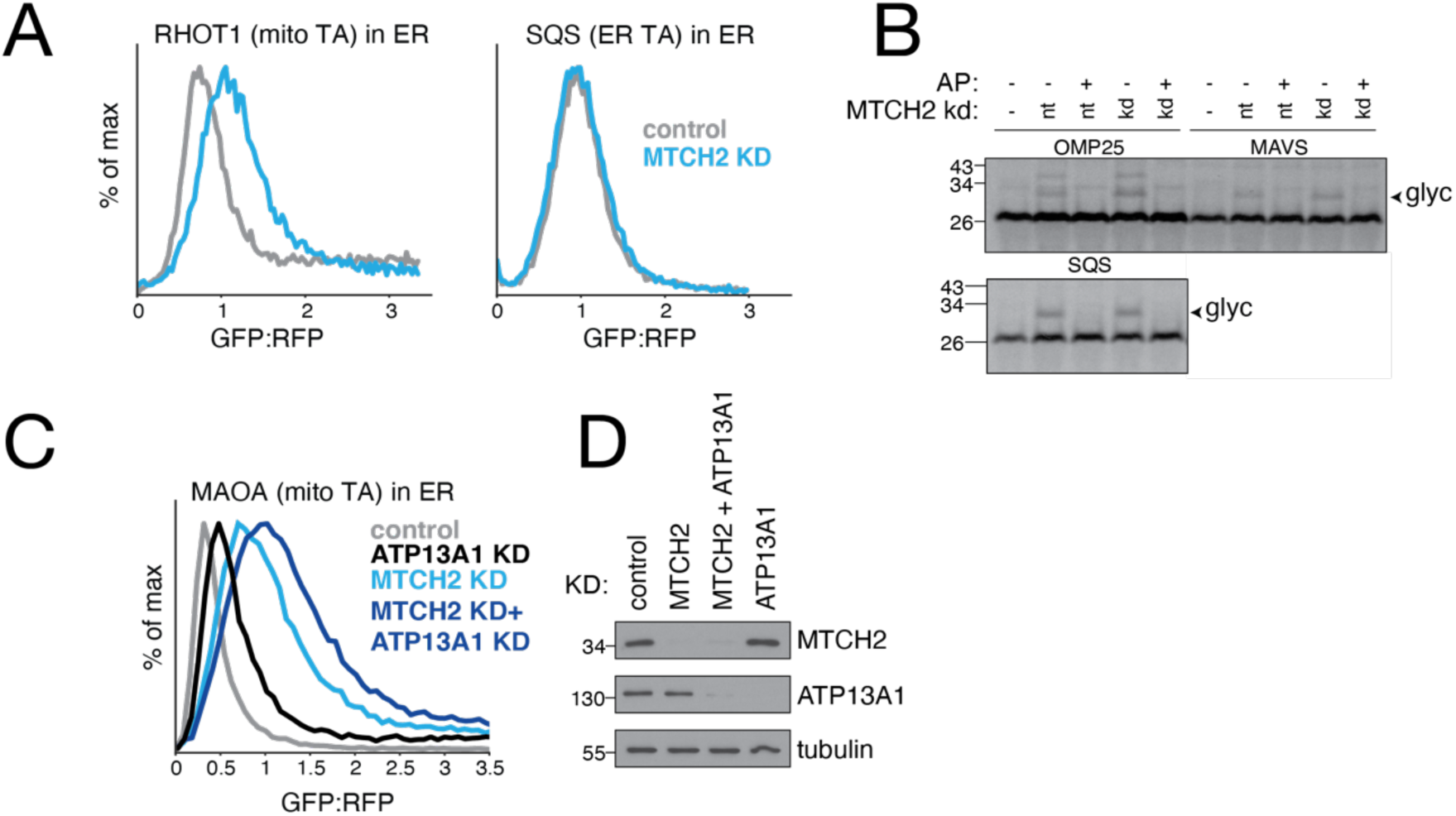
Depletion of MTCH2 causes increased mislocalization of mitochondrial TAs to the ER. **(A)** Cell lines expressing GFP1-10 in the ER lumen were used to monitor mislocalization to the ER of mitochondrial TAs fused to a C-terminal GFP11. Flow cytometry analysis of insertion into the ER of a mitochondrial TA (RHOT1) or an ER resident protein (SQS) in cells depleted of MTCH2. **(B)** Mislocalization of mitochondrial TAs in wildtype versus MTCH2 depleted cells was analyzed in vitro by appending a C-terminal opsin tag to their C-termini. The substrates were translation in reticulocyte lysate, puromycin treated and then mixed with semi-permeabilized cells. ER localization was detected as a glycosylated species (‘glyc’). Acceptor peptide (AP) was used to confirm the higher molecular weight band corresponded to a glycosylated species. Non-TA and ER resident TA controls were used to confirm the effect was specific to mislocalized mitochondrial TAs. **(C)** As in (A) but MTCH2 was depleted alongside the ER quality control factor ATP13A1. (D). Blots depicting MTCH2 and ATP13A1 levels in cells used for reporter assays in (C).

**Fig S17.**
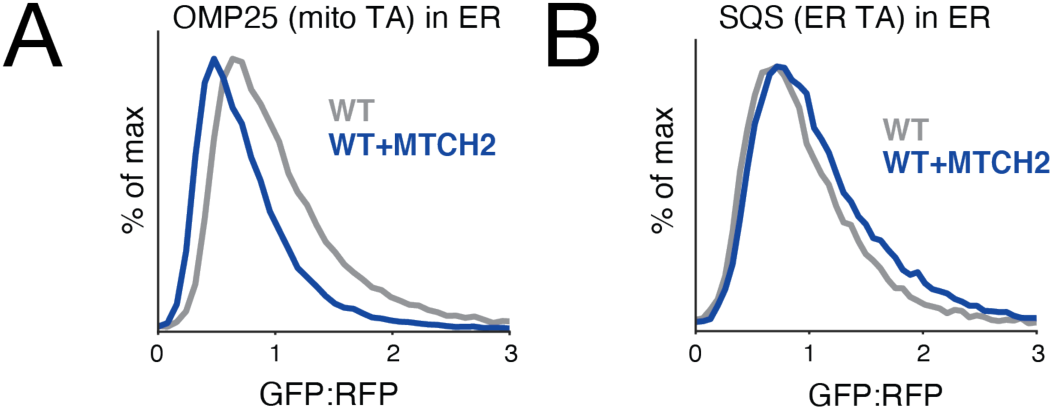
MTCH2 overexpression results in less mislocalization of mitochondrial TAs to the ER. As in fig. S16 but under conditions where MTCH2 is over-expressed. Note that there is a baseline of mislocalized OMP25-GFP11 reporter in the ER, likely due to over-expression of the construct.

