## Supplementary Materials for "MTCH2 is a mitochondrial outer membrane protein insertase"

Fig. S1

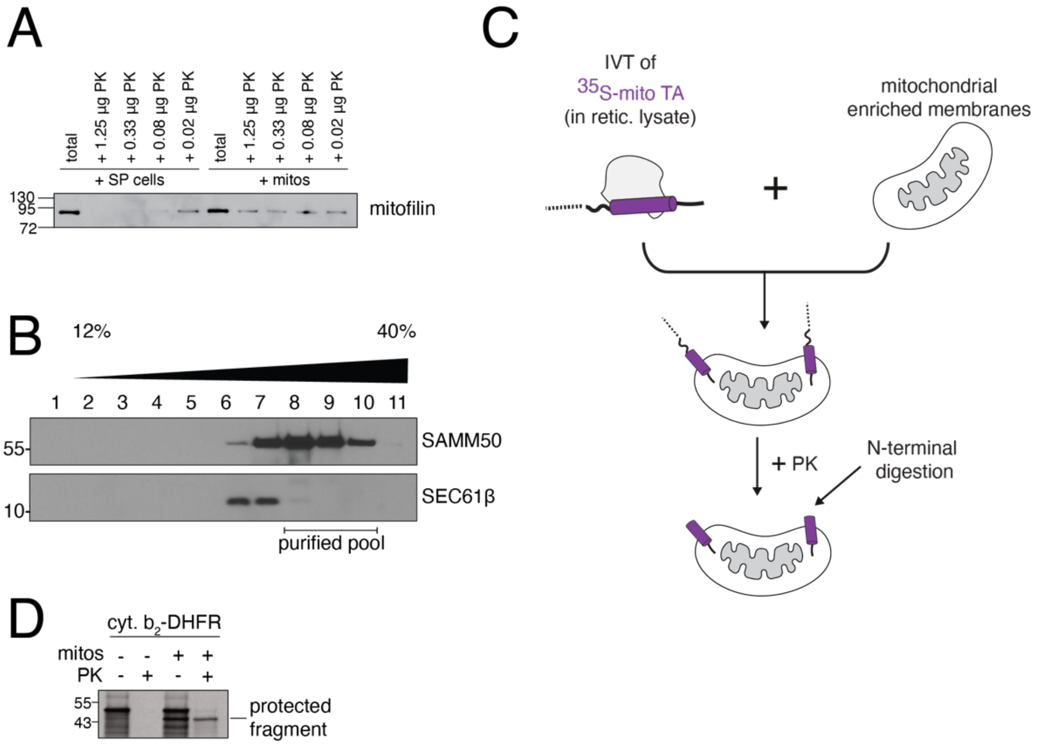


**Fig S1. In vitro protease protection assays into human mitochondria**. **(A)** The integrity of the outer membrane was tested using both semi-permeabilized K562 cells and mitochondrially enriched membrane fractions by treating each with decreasing amounts of proteinase K (PK). The resulting reactions were analyzed by western blot for the inner mitochondrial membrane protein mitofilin, which should be inaccessible to PK when the outer membrane is intact. **(B)** Further purification of mitochondria from K562 cells using a percoll gradient and density centrifugation. Blots for a mitochondrial marker (SAM50) and an ER marker (SEC61β) demonstrate relative proportion of mitochondria and ER in the 'mitochondrial enriched membranes' that are used for most in vitro experiments. Proteomics and crosslinking-mass spectrometry experiments were performed with percoll-gradient enriched mitochondria and used fractions 8-10 (Fig. 2A, Fig. 3C). **(C)** Schematic of the basic in vitro insertion assay, using protease protection as a readout for insertion. In all insertion assays, substrates were translated using an in vitro translation reaction (IVT) in rabbit reticulocyte lysate and $^{35}$S-methionine labelled, permitting detection by autoradiography. Translation was terminated by addition of puromycin, followed by incubation with membranes enriched in mitochondria derived from human K562 cells. After incubating with membranes, reactions were treated with PK. For TA proteins, the resulting protease protected band was immunoprecipitated via a tag on its C-terminus, ensuring insertion into the outer membrane in the correct orientation, and analyzed by SDS-PAGE and autoradiography. **(D)** The IMS-localized TOM substrate composed of the cytochrome b targeting sequence fused to the inert protein DHFR (cyt. $b_2$-DHFR) was translated and incubated with purified mitochondria to confirm their activity. Insertion was detected by protease protection as described.

Fig. S2

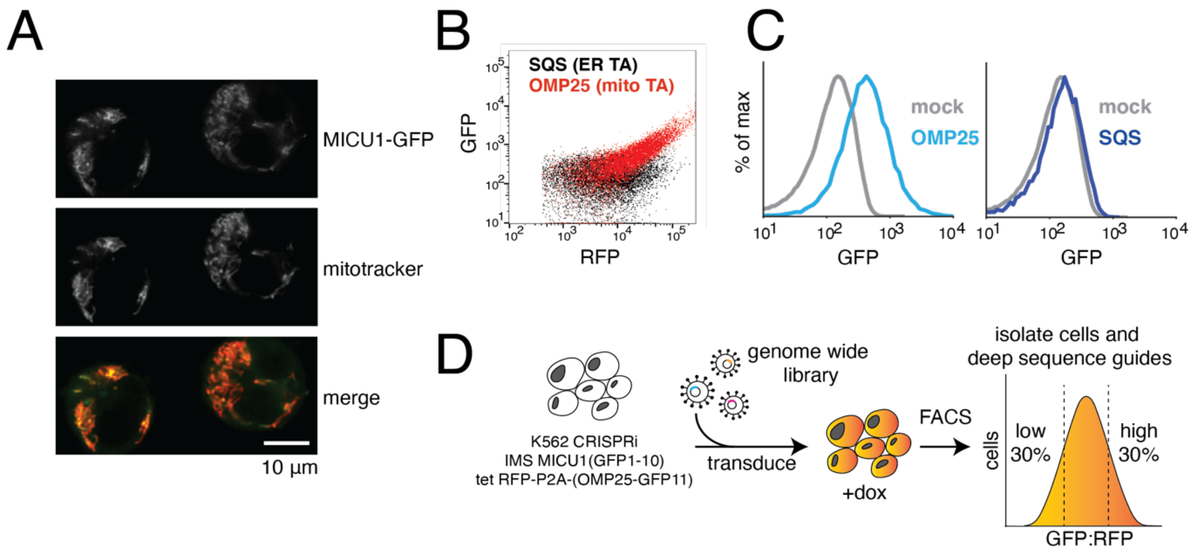


**Fig S2. A CRISPRi screening platform to identify factors involved in mitochondrial TA protein biogenesis in human cells**. **(A)** Microscopy showing that the IMS-targeting sequence from MICU1 conjugated to full length GFP results in its localization to the mitochondria in mammalian K562 CRISPRi cells. **(B)** The endogenous sequences of two TA proteins: SQS, which is localized to the ER, and OMP25, which under these conditions is dual-localized to both the outer membrane and ER, were appended to a C-terminal GFP11 in a backbone containing a translational control (RFP) separated by a viral 2A sequence (see Fig. 1B). These constructs were independently introduced into cells expressing IMS localized GFP1-10 and analyzed by flow cytometry. **(C)** Histograms of (B) comparing GFP fluorescence for OMP25 and SQS compared to a mock transduced control. The marked increase in GFP fluorescence suggest that OMP25, but not SQS can successfully conjugate with GFP1-10 localized to the IMS. **(D)** Workflow of the FACS-based CRISPRi screen. A K562 CRISPRi reporter cell line was constructed that constitutively expressed GFP1-10 in the IMS and the OMP25-GFP11 reporter under an inducible promoter. For the screen, these cells were transduced with a genome-scale CRISPRi sgRNA library and then the OMP25-GFP11 reporter was induced with doxycycline for 24 hours prior to cell sorting. Cells were sorted based on ratiometric changes in GFP relative to RFP, and sgRNAs expressed in the isolated cells were identified using deep sequencing.

Fig. S3

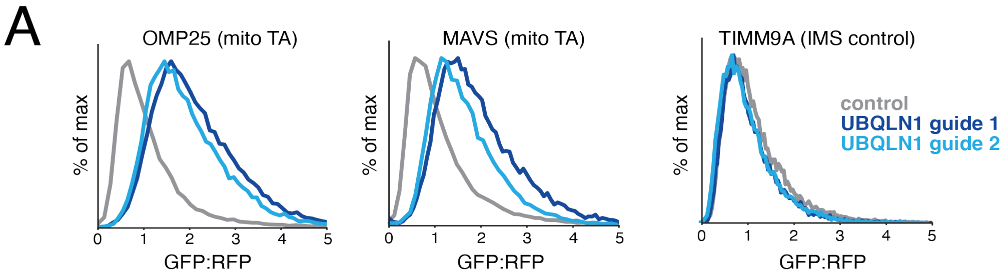


**Fig S3. UBQLN1 is a quality control factor for mitochondrial TAs**. K562 CRISPRi cells expressing IMS GFP1-10 were depleted of UBQLN1 using two different sgRNAs. Reporters were introduced for either two mitochondrial TAs (OMP25 and MAVS) or an IMS localized control (TIM9A) and cells were analyzed by flow cytometry. Lack of UBQLN results in a ratiometric increase in GFP:RFP fluorescence for mitochondrial TAs, consistent with its previously reported role in targeting mislocalized mitochondrial TAs for degradation by the ubiquitin proteasome pathway (*16*).

Fig. S4

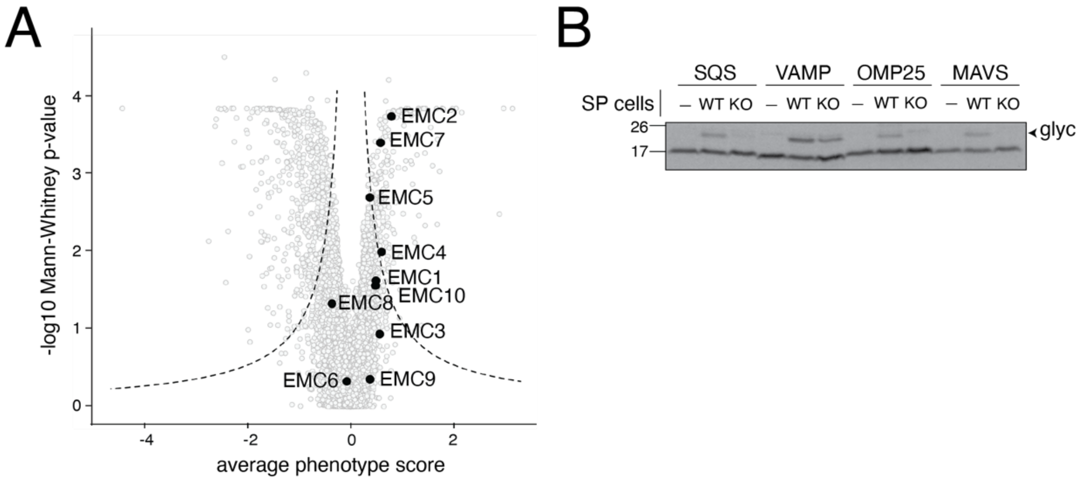


**Fig S4. The EMC is required for insertion of mislocalized mitochondrial TAs to the ER**. **(A)** Volcano plot of the genome-wide CRISPRi screen with EMC subunits shown in black. **(B)** A panel of ER (SQS and VAMP) and mitochondrial (OMP25 and MAVS) TA proteins were conjugated to a C-terminal opsin epitope. The opsin epitope contains a consensus glycosylation sequence that is modified upon insertion into the ER lumen. These constructs were then translated in rabbit reticulocyte lysate in the presence of $^{35}$S-methionine. The reactions were puromycin treated and incubated with either wild-type (WT) or EMC knockout (KO) semi-permeabilized (SP) cells. Insertion into the ER, as monitored by appearance of a glycosylated band (‘glyc’), was dependent on the EMC for its canonical substrate SQS, and both mitochondrial TAs. In contrast, VAMP’s insertion was unaffected by EMC knockout, consistent with its previously reported dependence on the GET pathway for insertion (*26*).

Fig. S5

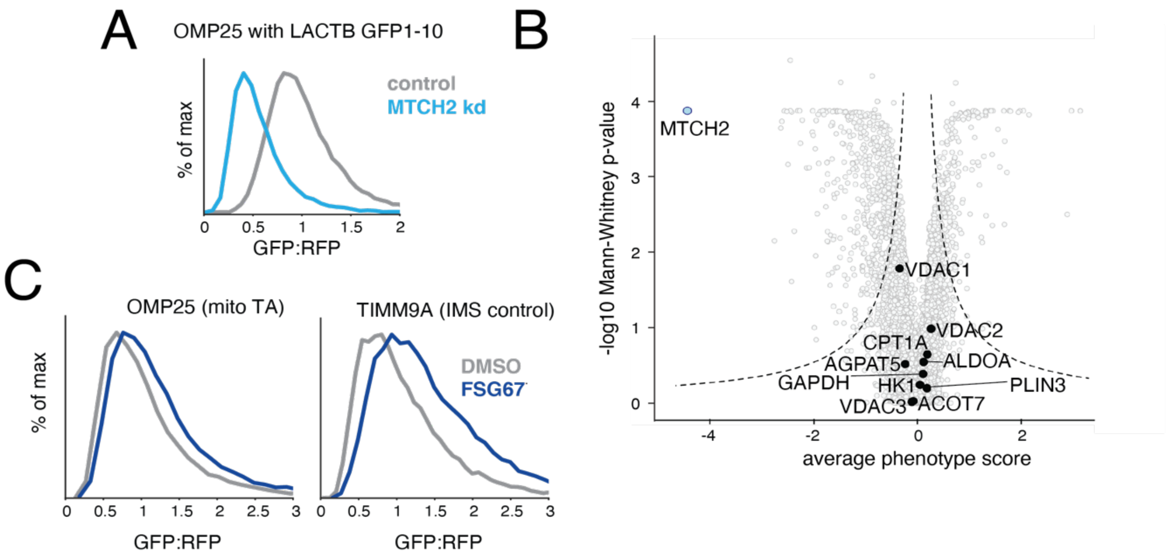


**Fig S5**. **Assessing the effects of lipid biogenesis defects on mitochondrial TAs. (A)** Flow cytometry analysis as in Fig. 1D but with an alternative IMS targeting sequence derived from LACTB appended to the GFP1-10 (*44*). (**B**) Volcano plot of the genome-wide CRISPRi screen indicating MTCH2 (in light blue) and factors previously implicated in the regulation of outer membrane fatty acid synthesis or transport (in black). **(C)** K562 IMS GFP1-10 expressing cells were treated with the pan GPAT inhibitor FSG67 for 16 hours (75 µM) or a vehicle. MTCH2-dependent mitochondrial fusion has been shown to require Glycerol 3-phosphate acyltransferases (GPATs) catalysed LPA synthesis. A reporter expressing either a mitochondrial TA (OMP25) or an IMS localized control (TIM9A) were expressed in GPATi cells and analyzed by flow cytometry.

Fig. S6

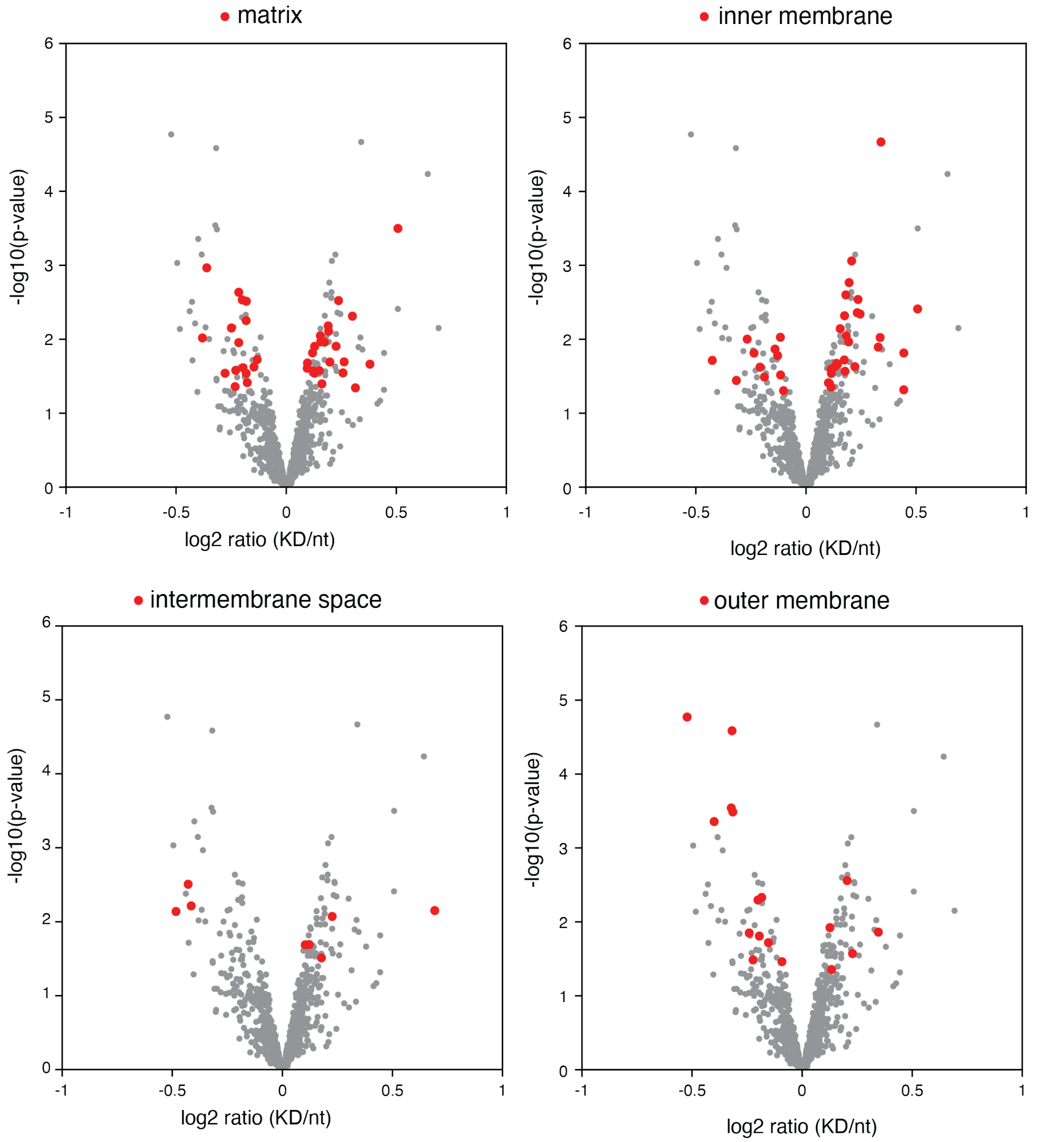


**Fig. S6**. **MTCH2 depletion affects endogenous outer mitochondrial membrane proteins.** As in Fig. 2A, but with all proteins from the indicated mitochondrial compartment which have a p-value > 0.05 highlighted in red.

Fig. S7

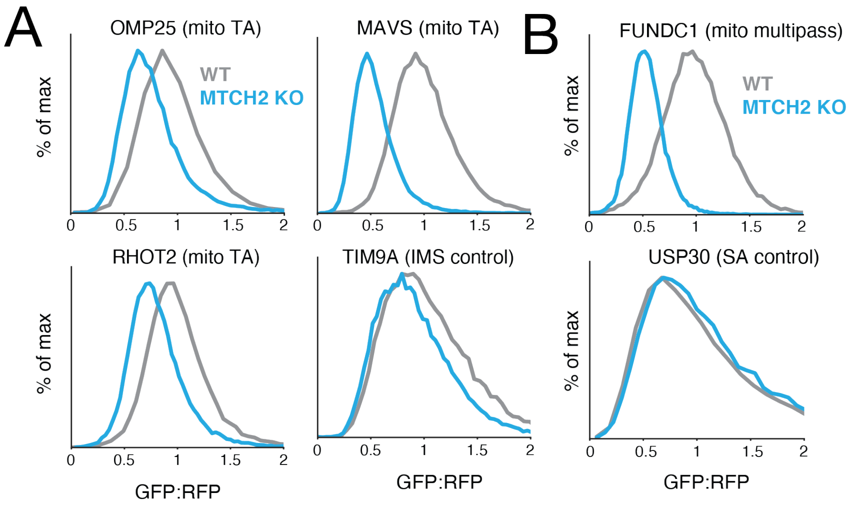


**Fig. S7**. **Analysis of TA proteins for MTCH2 dependent biogenesis in knockout cells.** As in Fig. 2C but in wild type compared to MTCH2 knockout cells. Histograms summarizing flow cytometry analysis of the integration of the indicated mitochondrial proteins.

Fig. S8

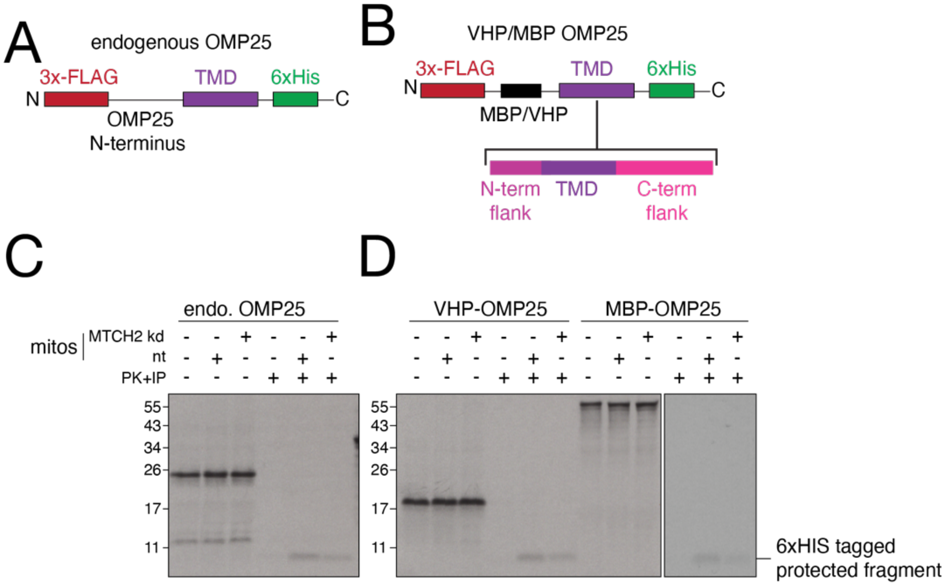


**Fig. S8. Establishing an in vitro system to test differential dependence on MTCH2 on insertion of a panel of mitochondrial TAs. (A, B)** Schematic of OMP25 constructs used to test whether the dependence on MTCH2 for insertion that we observed with the full-length endogenous OMP25 (A) could be recapitulated with an artificial N-terminus (B). **(C)**Using the protease protection assay described in fig. S1, we compared insertion of the endogenous OMP25 containing a C-terminal 6xHIS tag into mitochondria isolated from K562 cells expressing either a non-targeting (nt) or MTCH2 sgRNA (kd). We observed that loss of MTCH2 specifically decreased the levels of a protease protected fragment consistent in size with the TMD and C-terminus of OMP25 (compare lanes 5 vs 6). In the absence of mitochondria, no protected fragment is observed (lane 4). Because this protease protected fragment could be immunoprecipitated using a C-terminal 6xHIS tag, we verified that OMP25 was inserted in the correct orientation, with its C-terminus in the IMS and its N-terminus facing the cytosol. **(D)** As in (A) but using a fusion of the OMP25 TMD and its flanking residues to both the unrelated globular proteins MBP and VHP. We concluded that the OMP25 TMD alone is sufficient to confer MTCH2 dependent insertion on both of the tested fusion proteins (compare lanes 5 vs 6 and 11 vs 12). Because the VHP-fusions were translated more efficiently, we generated a panel of mitochondrial TAs using the depicted VHP N-terminus as shown in (B).

Fig. S9

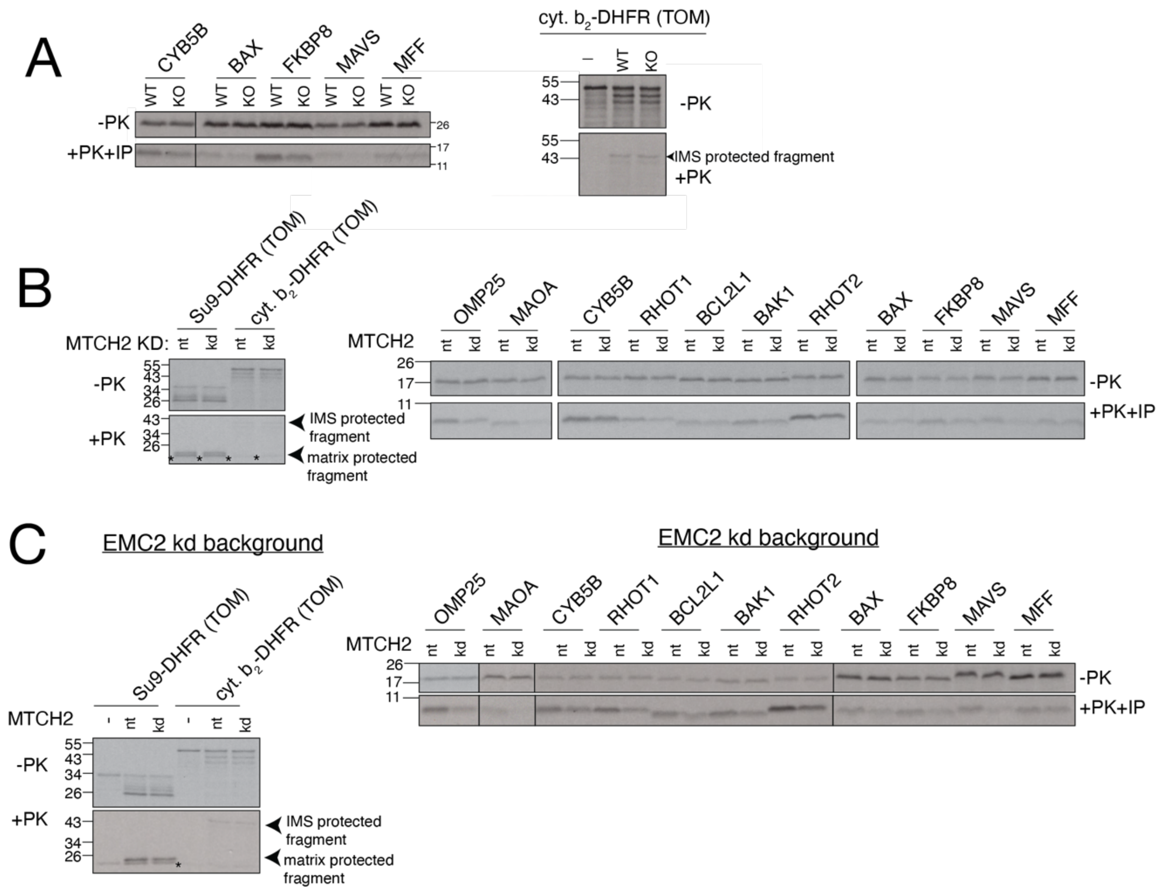


**Fig. S9. Insertion of mitochondrial TAs in vitro is affected by MTCH2 knockdown. (A)** Additional substrates were tested in parallel with the data displayed in Fig. 3A, where the indicated TMDs and flanking residues were fused to VHP and examined for insertion into wild type (WT) and MTCH2 knockout (KO) mitochondria. As a control, a canonical IMS localized TOM substrate derived from a fusion of the cytochrome b targeting sequence to DHFR (cyt. $b_2$-DHFR) was tested in parallel. **(B)** As in Fig 3A except using mitochondria isolated from K562 CRISPRi cells expressing either a non-targeting (nt) or MTCH2 targeting (kd) sgRNA. A panel of mitochondrial TAs was tested in parallel with a matrix (Su9-DHFR) and IMS (cyt. $b_2$-DHFR) targeted control that rely on the TOM pathway, which were unaffected**.** *Denotes the folded, protease resistant, DHFR domain that migrates immediately below the mature, matrix targeted control, and is visible in the absence of mitochondria. **(C)** Note that because some residual ER is present in the enriched mitochondrial membranes used in these insertion reactions (see fig. S1), for those substrates that are dual localized, we cannot formally differentiate between insertion into mitochondria vs ER in this assay alone. To address this, we have tested the complete TA panel in an EMC knockdown background, which we found eliminated mistargeting of mitochondrial TAs to the ER (fig. S4). Therefore, we perform insertion assays as in (A) except using (see fig. S1B) mitochondria isolated from either EMC2 knock down or EMC2 and MTCH2 knock down K562 CRISPRi cells. The insertion defect in the absence of MTCH2 was enhanced for some substrates (see CYB5B) when entry to the ER was blocked, suggesting these substrates may be partially dual localized under these conditions.

Fig. S10

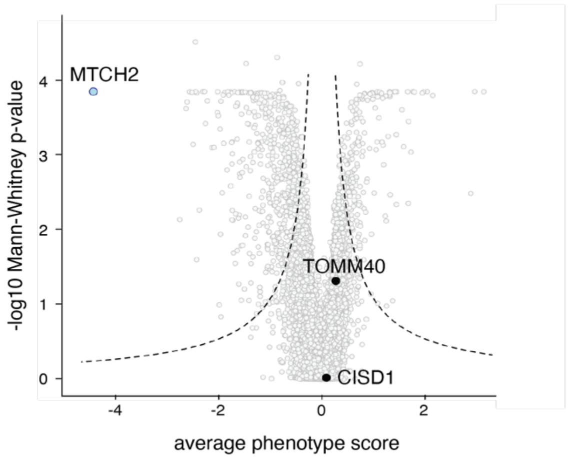


**Fig. S10. TOM40 and CISD1 are not significant hits in a mitochondrial TA biogenesis CRISPRi screen.** Volcano plot of the genome-wide CRISPRi screen as in Fig. 1C indicating MTCH2 (in light blue) and the two other factors (TOM40 and CISD1, in black) that were enriched upon crosslinking OMP25 with purified mitochondria (Fig. 3C; see fig. S1C for mitochondrial purification).

Fig. S11

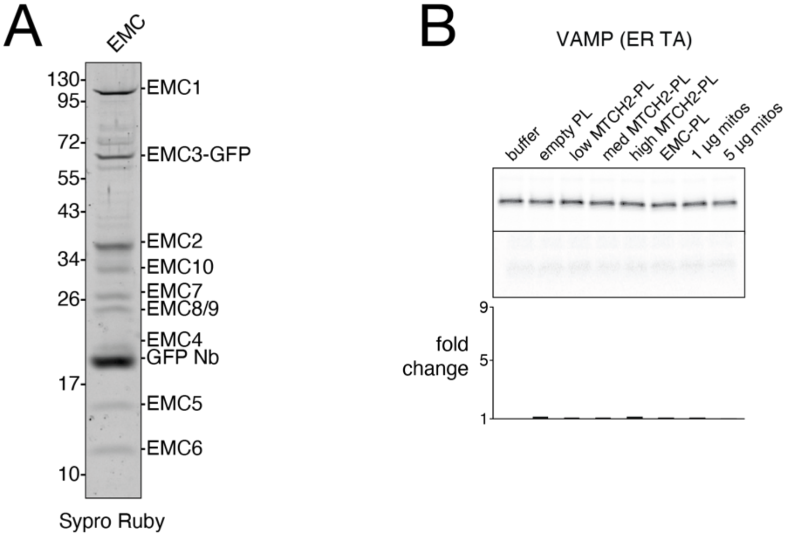


**Fig. S11. Purification of the human EMC and ER TA insertion control. (A)** The human EMC was expressed and purified as previously described using a GFP fused to the C-terminus of EMC3 in the detergent DBC (*25*) and visualized using Sypro Ruby staining. This was further used for generating EMC-proteoliposomes as indicated in Fig. 3E-F. (B) As in Fig. 3F but with an ER TA protein (VAMP) that is dependent on the GET-pathway for insertion.

Fig. S12

A

| alignment | | % ID |
|---|---|---|
| SLC25A46 | MTCH1 | 16.5 |
| SLC25A46 | MTCH2 | 10.7 |
| MTCH1 | MTCH2 | 36.4 |

B

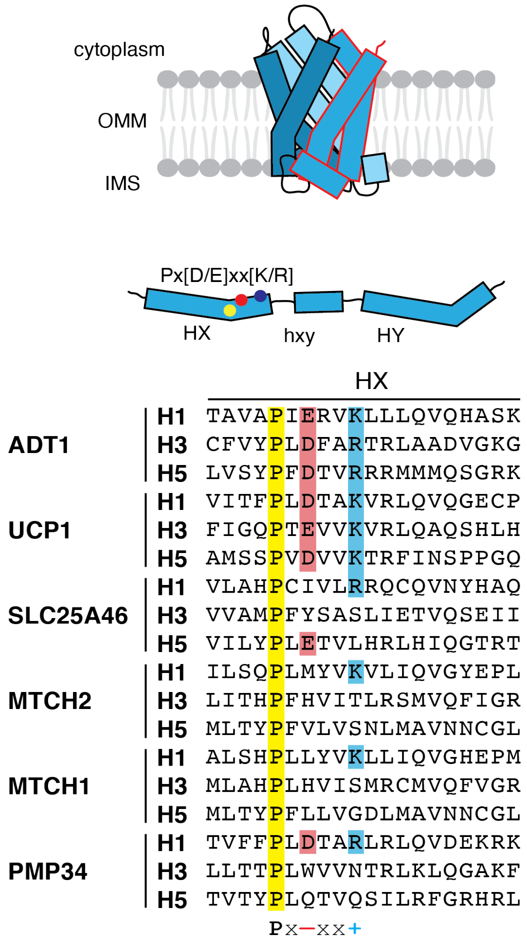


**Fig S12. Shared features of outer membrane SLC25 transporters**. **(A)** Sequence identity derived from pairwise alignment between SLC25A46, MTCH1, and MTCH2. **(B)** On top, a cartoon representation of SLC25 TMD arrangement showing the 3 SLC25 repeats in unique shades of blue, with a single repeat outlined in red. In the middle, a schematic showing the location of characteristic motifs within a single SLC25 repeat, which normally encodes 2 TM helices. On bottom, sequence alignment of all individual SLC25 repeats from 2 inner membrane SLC25 transporters (ADT1, UCP1) and 4 outer membrane SLC25 transporters, three mitochondrial (SLC25A46, MTCH1, MTCH2) and one peroxisomal (PMP34), with residues from the Px[D/E]xx[K/R] motif highlighted.

Fig. S13

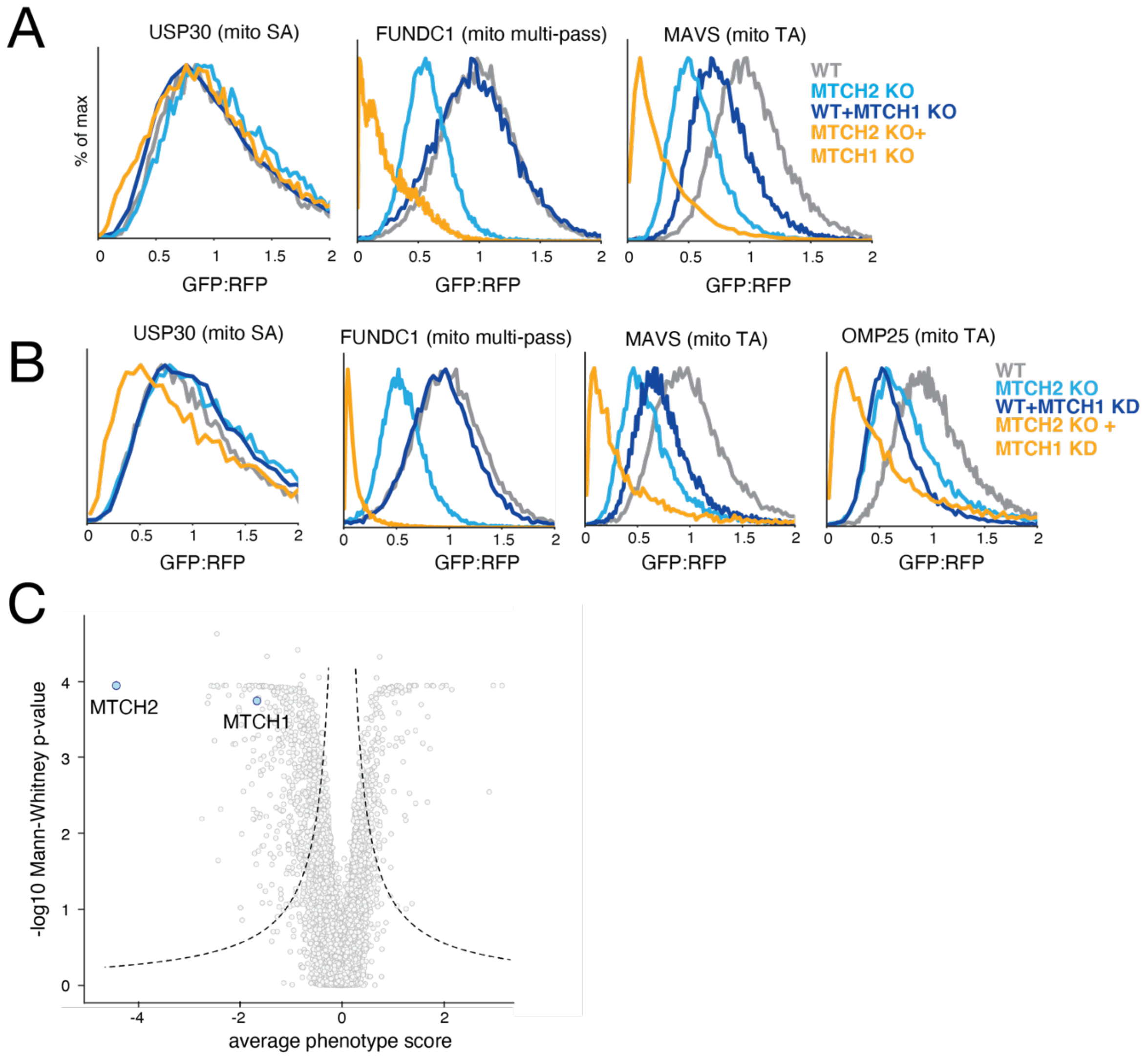


**Fig. S13. MTCH1 acts in a parallel pathway to mediate insertion of mitochondrial TAs. (A)** As in Fig. 4B for a set of outer membrane reporters including a signal-anchored protein (USP30), a mitochondrial TA (MAVS) and a multipass protein (FUNDC1). **(B)** As in (A) but in either wild-type or MTCH2 KO K562 CRISPRi cells expressing guides targeting MTCH1 for knock-down. **(C)** Volcano plot of the genome-wide CRISPRi screen with MTCH1 and MTCH2 highlighted.

Fig. S14

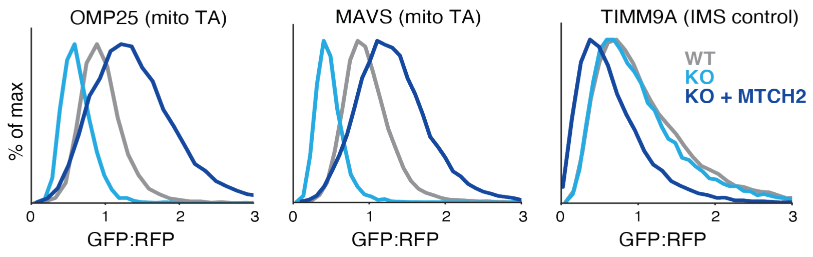


**Fig. S14. MTCH2 overexpression drives insertion of mitochondrial TAs into the outer membrane.** Flow cytometry analysis of two mitochondrial TAs (OMP25 and MAVS) and an IMS control (TIM9A) in the indicated cell types.

Fig. S15

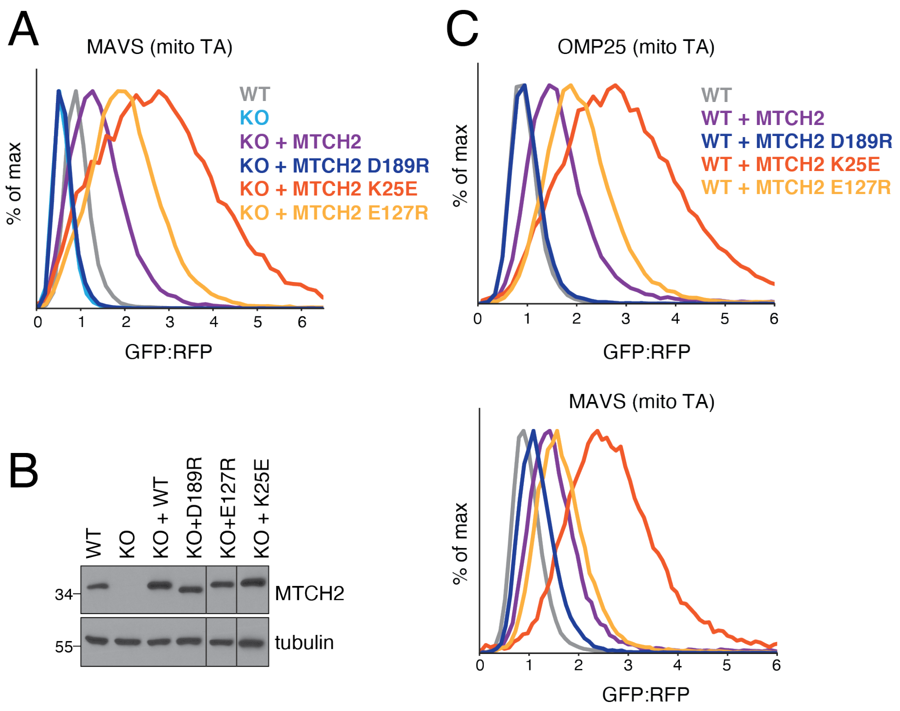


**Fig. S15. Point mutants to MTCH2 TMDs affect integration of mitochondrial TAs**. **(A)** As in Fig. 4C for another mitochondrial TA (MAVS). **(B)** Blots showing expression of wild-type MTCH2 and the indicated point mutants in a K562 MTCH2 KO background relative to wild-type cells. **(C)** As in (A) except point mutants were expressed on top of wild-type (WT) cells which are expressing functional copies of MTCH2.

Fig. S16

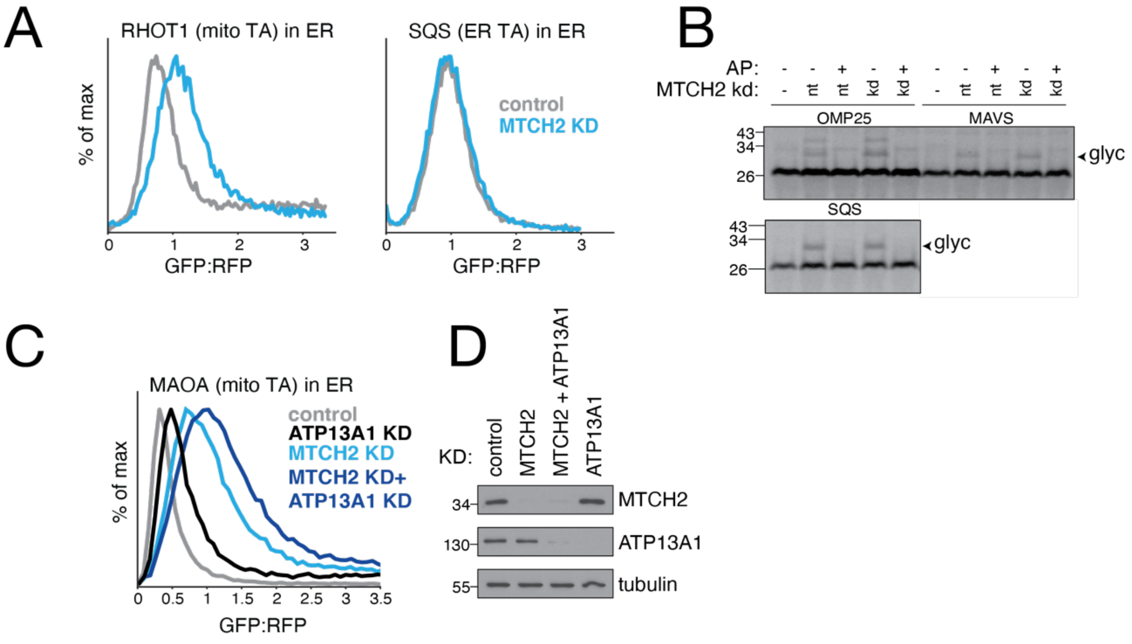


**Fig. S16. Depletion of MTCH2 causes increased mislocalization of mitochondrial TAs to the ER.** **(A)** Cell lines expressing GFP1-10 in the ER lumen were used to monitor mislocalization to the ER of mitochondrial TAs fused to a C-terminal GFP11. Flow cytometry analysis of insertion into the ER of a mitochondrial TA (RHOT1) or an ER resident protein (SQS) in cells depleted of MTCH2. **(B)** Mislocalization of mitochondrial TAs in wildtype versus MTCH2 depleted cells was analyzed in vitro by appending a C-terminal opsin tag to their C-termini. The substrates were translation in reticulocyte lysate, puromycin treated and then mixed with semi-permeabilized cells. ER localization was detected as a glycosylated species ('glyc'). Acceptor peptide (AP) was used to confirm the higher molecular weight band corresponded to a glycosylated species. Non-TA and ER resident TA controls were used to confirm the effect was specific to mislocalized mitochondrial TAs. **(C)** As in (A) but MTCH2 was depleted alongside the ER quality control factor ATP13A1. (D). Blots depicting MTCH2 and ATP13A1 levels in cells used for reporter assays in (C).

Fig. S17

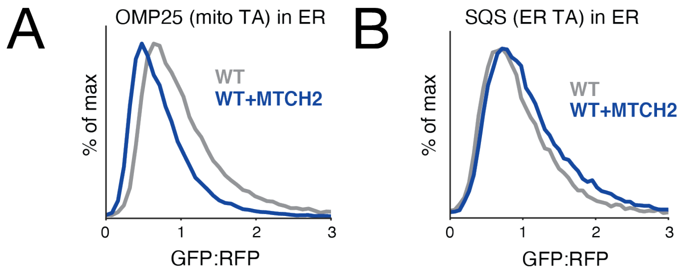


**Fig S17. MTCH2 overexpression results in less mislocalization of mitochondrial TAs to the ER.** As in fig. S16 but under conditions where MTCH2 is over-expressed. Note that there is a baseline of mislocalized OMP25-GFP11 reporter in the ER, likely due to over-expression of the construct.
